# Oxysterol-sensing by Liver X receptor counteracts ferroptosis via lipid remodeling

**DOI:** 10.64898/2026.08.17.745193

**Authors:** Andrea Kolak, Juliane Tschuck, Shibo Sun, Judith Sailer, Oliver Hartmann, Ina Rothenaigner, Huiyun Hu, Elizaveta Efanova, Fabien Riols, Mark Haid, Hans Zischka, Markus Diefenbacher, José Pedro Friedmann Angeli, Kamyar Hadian

## Abstract

Ferroptosis, an iron-dependent form of regulated cell death, is controlled by cellular metabolism. Nutrients and metabolites determine cell states that render cells sensitive or resistant to ferroptosis. Nuclear receptors can act as cellular sensors for distinct metabolites and nutrients to regulate ferroptosis. We performed a chemical genetics screen using a nuclear receptor small molecule library to identify novel regulators of ferroptosis. We find that activating or overexpressing the liver X receptor (LXR) suppresses ferroptosis in various cell models, including *ex vivo* primary mouse hepatocytes. Interestingly, hepatocellular carcinoma with high levels of LXR shows poorer survival outcomes. In cells, activation of LXR by the endogenous oxysterol 24(S),25-epoxycholesterol or synthetic agonists reduces lipid peroxidation and ferroptotic cell death. Mechanistically, LXR activation drives a selective transcriptional program upregulating SREBP-1c, SCD1 and ACSL3, key enzymes involved in the synthesis of monounsaturated fatty acid-containing phospholipids (MUFA-PLs). Lipidomic analysis reveals that this lipid remodeling enriches cellular membranes with MUFA-PLs, reducing their susceptibility to peroxidation and thereby counteracting ferroptosis. Together, we identify LXR as an oxysterol-sensing endogenous suppressor of ferroptosis coupling oxysterol sensing to the adaptive remodeling of cellular membrane lipid composition to limit lipid peroxidation.

## Introduction

Both cell division and cell death are essential to maintaining normal tissue homeostasis in multicellular organisms. Ferroptosis, a regulated form of cell death, is distinct from other cell death modalities such as apoptosis, because it is metabolically driven rather than induced by a cascade of signaling events^1, 2^. It is characterized by iron-dependent lipid peroxidation^3^. The incorporation of polyunsaturated fatty acids (PUFAs) into phospholipids is facilitated by the actions of acyl-CoA synthetase long-chain family member 4 (ACSL4) and lysophosphatidylcholine acyltransferase 3 (LPCAT3)^2, 4^. To inhibit ferroptosis, cells deploy multiple defense strategies that limit lipid peroxidation. The system x_c_^−^–glutathione–GPX4 axis is the primary ferroptosis suppression pathway^3^. Additionally, two GPX4-independent ferroptosis defense systems have been identified: the FSP1–ubiquinol–vitamin K axis^5–7^ and the GCH1–DHFR–tetrahydrobiopterin axis^8, 9^. Another mechanism that counteracts ferroptosis is the shift toward a membrane rich in monounsaturated fatty acids (MUFAs), catalyzed by acyl-CoA synthetase long-chain family member 3 (ACSL3)^10^ and membrane-bound O-acyltransferase domain-containing 1/2 (MBOAT1/2)^11^.

Several transcription factors have been identified as key regulators of ferroptosis, including NRF2, ATF4, and YAP1^12^. NRF2 acts as a central antioxidant guardian by transcriptionally activating genes involved in thioredoxin and glutathione metabolism, as well as the primary ferroptosis suppressors SLC7A11 and genes important for glutathione synthesis^13^. Similarly, ATF4 increases SLC7A11 expression, thereby expanding intracellular glutathione reserves, which are necessary for GPX4 activity and ferroptosis resistance^12^. Nuclear receptors (NRs) are a class of transcription factors activated by various metabolites, including fatty acids, bile acids, vitamins, hormones, and oxysterols. NRs are structurally defined by ligand-binding and DNA-binding domains, and they typically function as heterodimers to coordinate comprehensive transcriptional programs^14^. Several nuclear receptors have been shown to suppress ferroptosis, these include the fatty acid-sensitive nuclear receptor peroxisome proliferator-activated receptor alpha (PPARα)^15, 16^, the estrogen receptor (ER)^11, 17^, the farnesoid X receptor (FXR)^18, 19^ and the retinoic acid receptor (RAR)^20^. Together, these studies underline that nuclear receptors play a key role in sensing metabolites, lipids, hormones or vitamins to control ferroptosis.

In this study, screening a chemical library of nuclear receptor agonists and antagonists revealed that LXR activation suppresses ferroptosis. As LXR is a sensor for oxysterols, some of which arise from ROS-mediated oxidation of cholesterol under oxidative conditions resembling those of ferroptotic lipid peroxidation, we reasoned that LXR may sense oxidative lipid stress and counteract it. We demonstrate that LXR activation increases the expression of SREBP-1c, SCD1 and ACSL3, thereby promoting the incorporation of MUFAs into membranes and suppresses lipid peroxidation and ferroptosis. In this way, transcription serves as the means by which LXR couples oxysterol sensing to a protective membrane response.

## Results

### LXR is an endogenous oxysterol-sensing suppressor of ferroptosis

To systematically analyze the class of nuclear receptors (NR) regarding their influence on ferroptosis, we performed a compound screen using a library of 550 agonists or antagonists of various nuclear receptors. HT-1080 cells were treated with distinct NR agonists and antagonists of nuclear receptor library and then challenged with RSL3 at IC_80_ to induce ferroptosis (Fig. 1a). The potential of specific NR agonists or antagonists to inhibit ferroptosis was detected in a viability assay. The screen yielded several agonists of nuclear receptors previously reported as ferroptosis regulators, demonstrating the robustness of the screen. These included agonists of the ER^11, 17^, androgen receptor (AR)^11^, and FXR^18, 19^. Interestingly, another hit was GSK3987 (hereafter referred to as LXRag1), an agonist of the liver X receptor (LXR), a nuclear receptor, which has not yet been fully studied in the context of ferroptosis regulation (Fig. 1b). An analysis of over 800 cancer cell lines in the CTRP database^21^ revealed that cancer cells become more resistant to GPX4-targeting drugs (RSL3, ML162, and ML210) when the NR1H3 (LXRα) gene is overexpressed (Fig. 1c), confirming an association between LXR and ferroptosis. LXR is present in two isoforms, LXRα (NR1H3) and LXRβ (NR1H2), with high similarity. LXRα is largely expressed in the liver and other metabolic active tissues, while LXRβ is ubiquitously expressed^22^.

**Fig. 1.**
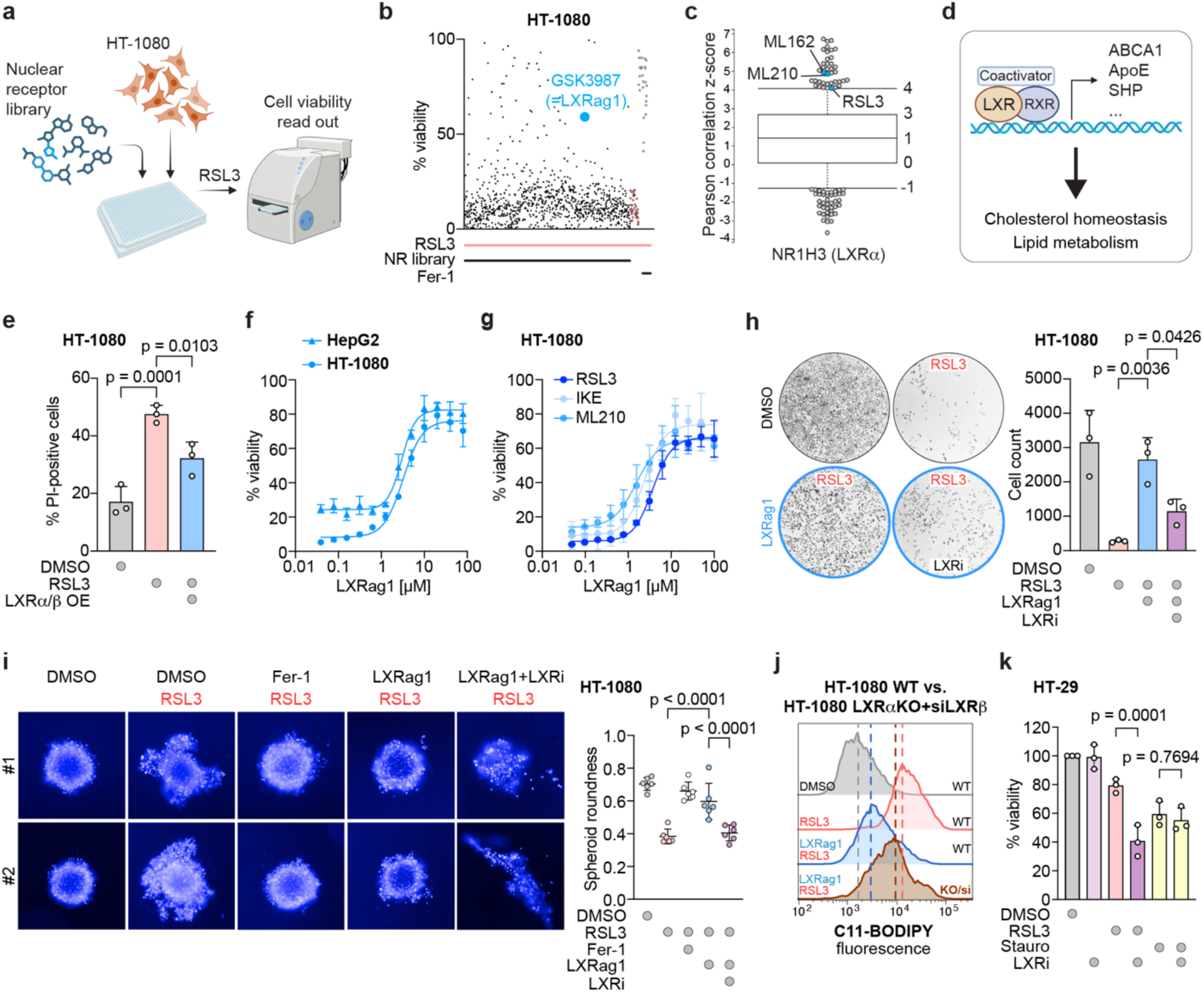
LXR activation protects from ferroptosis. **a,** Schematic overview of the ferroptosis screen performed using the nuclear receptor library on RSL3-treated HT-1080 cells. **b**, Scatter plot of the viability receptor screen in HT-1080 cells treated with 200 nM RSL3 (IC_80_) identified LXR agonist 1 (LXRag1) as a ferroptosis inhibitor. Ferrostatin-1 (Fer-1) was used as a positive control for ferroptosis inhibition. **c,** Positive Pearson correlation between high expression of NR1H3 (LXRα) and resistance to GPX4 inhibitors RSL3, ML210 and ML162 in cancer cells. **d**, Illustration of LXR activation leading to the transcription of target genes regulating cholesterol homeostasis and lipid metabolism. **e**, Cell death analysis by quantification of PI positive cells upon induction of ferroptosis with 200 nM RSL3. LXR-overexpressing HT-1080 cells show reduced cell death compared to control-transfected cells; mean ± SD of *n* = 3 independent biological replicates; ordinary one-way ANOVA with Šidák’s multiple-comparison test. **f**, LXR-activation dose-dependently inhibits ferroptotic cell death induced by 150 nM RSL3 in HT-1080 and HepG2 cells; mean ± SD of *n* = 3 independent biological replicates. **g**, LXR activation dose-dependently inhibits ferroptosis induced by RSL3 (150 nM), IKE (2 µM) and ML210 (300 nM) in HT-1080 cells; mean ± SD of *n* = 3 independent biological replicates. **h**, Crystal violet staining of HT-1080 cells treated with 150 nM RSL3 show increased cell density upon addition of LXRag1 (10 µM), which is reduced when LXR is simultaneously inhibited with LXRi (25 µM). Quantified cell counts measured are shown in the diagram, mean ± SD of *n* = 3 independent biological replicates; ordinary one-way ANOVA with Šidák’s multiple-comparison test. **i**, Spheroids of HT-1080 cells are disrupted upon treatment with 200 nM RSL3. LXR activation (10 µM LXRag1) protects spheroids from RSL3-induced disintegration, which is reversed by LXRi (25 µM). Fer-1 (2 µM) was used as positive control. Analysis of spheroid roundness is depicted in the diagram; mean ± SD of *n* = 8 independent biological replicates; ordinary one-way ANOVA with Šidák’s multiple-comparison test. **j**, Analysis of C11-BODIPY fluorescence shift indicates that activation of LXR (10 µM LXRag1) decreases lipid peroxidation in ferroptotic (1 µM RSL3) wild-type HT-1080 cells. This reduction is attenuated in 1 µM RSL3-treated LXRα-knockout+siLXRβ-transfected cells; *n* = 3 independent biological replicates. **k**, LXR inhibition with 25 µM LXR-Inhibitor (LXRi) enhances RSL3-induced loss of viability (1 µM RSL3) but does not affect viability alone or in combination with 1 µM staurosporine (Stauro) in ferroptosis resistant HT-29 cells; mean ± SD of *n* = 3 independent biological replicates; ordinary one-way ANOVA with Šidák’s multiple-comparison test.

LXR senses oxysterols, *e.g.,* 24(*S*),25-epoxycholesterol, to get activated. Oxysterols are oxygenated cholesterol forms, which are generated through oxidation by radicals or reactive oxygen species (ROS), but can also be a product of enzymatic reaction^23^. As ROS-driven cholesterol oxidation is a feature of a similar oxidative environment that drives ferroptosis, this places LXR as a potential sensor of oxidative lipid stress. LXR forms a heterodimeric complex with the retinoic X receptor (RXR) and upon activation initiates transcription of genes involved in cholesterol homeostasis and lipid metabolism^22^ (Fig. 1d).

Next, we overexpressed LXRα/β in HT-1080 cells to demonstrate that excess LXR has anti-ferroptotic capacity. Using propidium iodide (PI) to stain dead cells, we see that RSL3 increases the number of PI-positive cells, which is significantly reduced in LXRα/β overexpressing cells (Fig. 1e and Extended Data Fig. 1a, b). Importantly, we could see the ferroptosis protective effect of LXR activation by LXRag1 in HT-1080 as well as HepG2 cells (Fig. 1f), and against several ferroptosis inducers, such as GPX4 inhibitors (RSL3 and ML210) or the system xc-inhibitor IKE (Fig. 1g). However, LXRag1 did not inhibit other regulated cell death pathways such as apoptosis (Extended Data Fig. 1c) or necroptosis (Extended Data Fig. 1d). These findings were confirmed via crystal violet staining in HT-1080 cells, which showed that the LXR inhibitor GSK2033 (hereafter referred to as LXRi) can neutralize the inhibitory effect of LXRag1 on RSL3-induced ferroptosis (Fig. 1h). Finally, we validated these results in a 3D cell culture model of HT-1080. While RSL3 treatment led to the dissociation of HT-1080 spheroids, LXRag1 preserved their structural integrity, an effect that was again abrogated by LXRi (Fig. 1i). While CTRP data and LXR overexpression experiments established a protective role for LXR in ferroptosis regulation, we aimed to ensure that the small molecule LXRag1 does not produce off-target effects contributing to ferroptosis inhibition. To this end, we generated an HT-1080-LXRαKO cell line and transfected this line with siRNAs against LXRβ. Notably, we could not obtain a full LXRα knockout, and the siLXRβ reduced LXRβ levels by only 60-70% (Extended Data Fig. 2a). It seems that the cells cannot tolerate full loss of LXR. Using non-transfected HT-1080 cells, we induced lipid peroxidation by adding RSL3 and this was markedly reduced by LXRag1-mediated activation of LXR (Fig. 1j). In contrast, reduction in LXRα and LXRβ levels in HT-1080-LXRαKO+siLXRβ cells showed increased lipid peroxidation in the presence of LXRag1, which was comparable to RSL3-treated HT-1080 WT cells (Fig. 1j and Extended Data Fig. 2b and 2c). This shows that the anti-ferroptotic effect of LXRag1 is directed through the nuclear receptor LXR, but not by any off-target chemical properties. In addition, we used a cell-free BODIPY assay^19, 20, 24^ to directly test antioxidant activity of LXRag1. LXRag1 only marginally reduced C11-BODIPY oxidation by AAPH (Extended Data Fig. 1e), again indicating that its anti-ferroptotic effect is not attributed to general antioxidant properties. To further strengthen our findings that LXR regulates ferroptosis, we inhibited LXR in ferroptosis-resistant HT-29 cells to sensitize these cells towards ferroptosis. While treatment with LXRi alone did not affect cell viability and RSL3 caused only a 20% reduction in cell viability, their combination synergized to induce 60% cell death (Fig. 1k). These results demonstrate that LXR inhibition sensitizes HT-29 cells to ferroptosis induction. Notably, LXRi did not synergize with staurosporine to induce apoptosis (Fig. 1k).

**Fig. 2.**
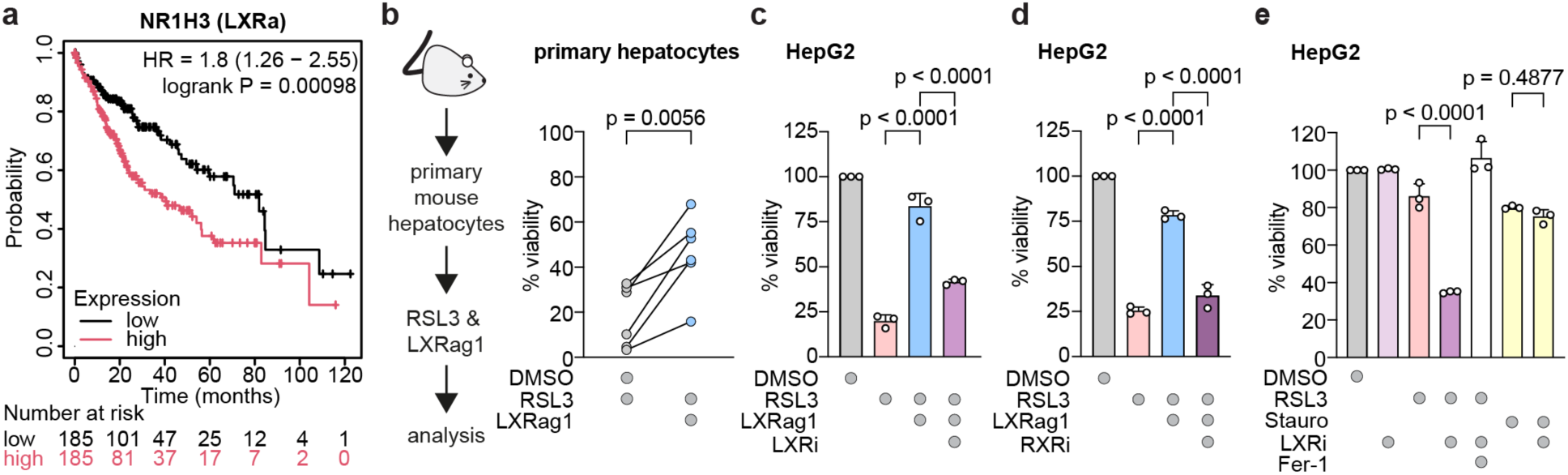
LXR/RXR transcriptional activity drives resistance to ferroptosis in liver cancer. **a,** Kaplan-Meier (KM) plot shows high levels of LXRα in liver hepatocellular carcinoma leads to worse patient survival rates **b,** Primary hepatocytes isolated from Hfe mice and treated with 200 nM RSL3 showed improved viability upon co-treatment with 10 µM LXRag1; mean ± SD of *n* = 6 independent biological replicates; ordinary one-way ANOVA with Šidák’s multiple-comparison test. **c,** Activation of LXR with 10 µM LXRag1 mitigates ferroptosis induced by 150 nM RSL3, whereas co-treatment with 25 µM LXRi attenuates this protective effect in HepG2 cells; mean ± SD of *n* = 3 independent biological replicates; ordinary one-way ANOVA with Šidák’s multiple-comparison test. **d**, Activation of LXR with 10 µM LXRag1 mitigates ferroptosis induced by 150 nM RSL3, whereas co-treatment with 25 µM RXR-Inhibitor (RXRi) attenuates this protective effect in HepG2 cells; mean ± SD of *n* = 3 independent biological replicates; ordinary one-way ANOVA with Šidák’s multiple-comparison test. **e**, LXR inhibition (25 µM LXRi) increases sensitivity of HepG2 cells to ferroptosis induced by a sublethal concentration of 50 nM RSL3 but does not alter viability alone or in combination with 1 µM staurosporine (Stauro). Fer-1 (2 µM) was used as positive control; mean ± SD of *n* = 3 independent biological replicates; ordinary one-way ANOVA with Šidák’s multiple-comparison test.

### LXR activity protects liver cells from ferroptosis and correlates with poor HCC prognosis

As LXR is well expressed and active in liver cells, we evaluated the clinical significance of LXRα in hepatocellular carcinoma (HCC). Therefore, we first examined its association with patient survival. A query of the KM Plotter^25–27^ revealed a negative correlation between LXRα expression in hepatocellular carcinoma and patient survival, with elevated LXRα levels being associated with poor prognosis (Fig. 2a). To further examine the role of LXR in liver cells, we extracted *ex vivo* primary mouse hepatocytes and treated them with RSL3 to induce ferroptosis. LXRag1 also attenuated RSL3-induced ferroptosis in primary mouse hepatocytes (Fig. 2b). Next, we treated the human liver cancer cell line, HepG2, with RSL3 and then added LXRag1, which again reduced ferroptosis. Notably, the combination of LXRag1 and LXRi treatments resensitized HepG2 cells to ferroptosis induction (Fig. 2c). A similar finding was obtained with RXR inhibitor (RXRi) treatment. Here, inhibiting the LXR-co-activator RXR with RXRi increased RSL3-induced cell death, even in the presence of LXRag1 (Fig. 2d). These results further demonstrate that the anti-ferroptotic activity of LXRag1 is mediated by LXR/RXR activity. To test the hypothesis that LXR inhibition may sensitize liver cells to ferroptosis as a therapeutic concept, we tested LXRi treatment in combination with a sublethal dose of RSL3. LXR inhibition alone did not induce cell death. However, LXRi increased the efficacy of a sublethal dose of RSL3, boosting it from 10% to 70% ferroptosis induction. This increase was blocked by Fer-1 (Fig. 2e). Again, LXRi had no effect on staurosporine-induced apoptosis (Fig. 2e), demonstrating a ferroptosis-specific effect.

Using various cell lines, spheroids, and *ex vivo* primary mouse hepatocytes alongside different chemical inducers of ferroptosis, we demonstrate that LXR activation or overexpression protects against ferroptotic cell death, while LXR inhibition increases sensitivity to ferroptosis.

### LXR activation reduces lipid peroxidation

A major hallmark of ferroptosis is the peroxidation of polyunsaturated fatty acyl tails (PUFAs). Therefore, we live-stained HepG2 (Fig. 3a) and HT-1080 (Fig. 3c) cells using the fluorescent lipid peroxidation sensor C11-BODIPY. In both cell lines, LXRag1 significantly reduced RSL3-induced BODIPY fluorescence, indicating that LXR activation suppresses lipid peroxidation in ferroptosis-sensitive cells. To validate this finding, we additionally immunostained 4-hydroxynonenal (4-HNE), a product of lipid peroxidation. LXR activation significantly reduced 4-HNE levels upon ferroptosis induction in HepG2 (Fig. 3b) and HT-1080 cells (Fig. 3d). Next, we performed the reciprocal experiment to induce ferroptosis in a ferroptosis-resistant cell line. Therefore, we used the ferroptosis-resistant HT-29 cells and observed that inhibition of LXR (LXRi) led to increased lipid peroxidation in combination with RSL3 treatment, as measured by C11-BODIPY levels (Fig. 3e).

**Fig. 3.**
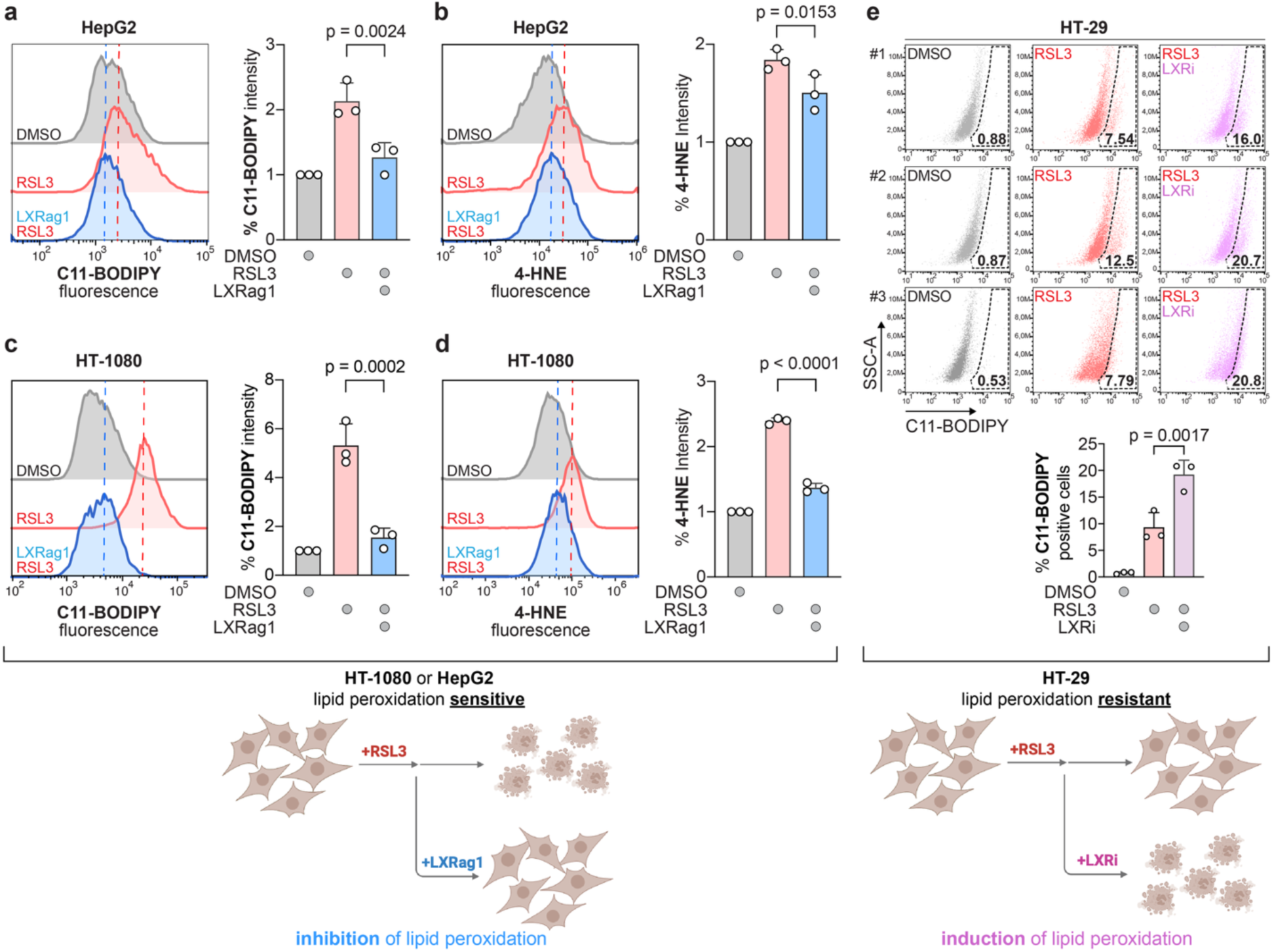
LXR activity negatively regulates lipid peroxidation. **a,** Fluorescence shift of C11-BODIPY indicates that activation of LXR (10 µM LXRag1) decreases lipid peroxidation in ferroptotic (1 µM RSL3) HepG2 cells; mean ± SD of *n* = 3 independent biological replicates; ordinary one-way ANOVA with Šidák’s multiple-comparison test. **b,** Reduced levels of 4-HNE measured in LXRag1-treated HepG2 cells compared to control cells treated with 1 µM RSL3 respectively; mean ± SD of *n* = 3 independent biological replicates; ordinary one-way ANOVA with Šidák’s multiple-comparison test. **c,** Fluorescence shift of C11-BODIPY indicates that activation of LXR (10 µM LXRag1) decreases lipid peroxidation in ferroptotic (200 nM RSL3) HT-1080 cells; mean ± SD of *n* = 3 independent biological replicates; ordinary one-way ANOVA with Šidák’s multiple-comparison test. **d**, Reduced levels of 4-HNE measured in LXRag1-treated HT-1080 cells compared to control cells treated with 200 nM RSL3 respectively; mean ± SD of *n* = 3 independent biological replicates; ordinary one-way ANOVA with Šidák’s multiple-comparison test. **e**, Increased amount of C11-BODIPY positive cells in LXRi (25 µM) and RSL3 (1 µM) treated HT-29 cells indicates an increase of lipid peroxidation upon LXR inhibition; mean ± SD of *n* = 3 independent biological replicates; ordinary one-way ANOVA with Šidák’s multiple-comparison test.

Together, our data show that modulation of LXR activity has an impact on the level of lipid peroxidation. Here, LXR activation suppresses ferroptotic cell death in lipid peroxidation sensitive cells, while LXR inhibition sensitizes toward ferroptosis in lipid peroxidation resistant cells.

### LXR selectively induces MUFA-PL synthesis genes

LXR is a nuclear receptor regulating the expression of genes involved in lipid metabolism and cholesterol homeostasis. Therefore, we investigated whether the anti-ferroptotic effect of LXR is mediated by transcriptional regulation of ferroptosis-regulating target genes. We performed qRT-PCR for a number of genes implicated in ferroptosis regulation. First, we checked receptor activation in HepG2 cells (Fig. 4a, Extended Data Fig. 3a and 3b). LXR activation by LXRag1 resulted in mRNA upregulation of LXRα and LXRβ, as well as the canonical target genes ApoE and SHP (Extended Data Fig. 3a). Interestingly, expressions of GPX4 and FSP1 were barely changed, while GCH1 expression was moderately upregulated (Extended Data Fig. 3b). This finding showed a clear distinction from FXR-activated ferroptosis suppression^19^. Instead, analysis of SREBP-1c, SCD1 and ACSL3 in HepG2 demonstrated marked upregulation upon LXR activation (Fig. 4a). Co-treatment with LXR inhibitor reduced SREBP-1c, SCD1 and ACSL3 expression (Fig. 4a), proving that this activation is nuclear receptor mediated. SREBP-1c, SCD1 and ACSL3 are key regulators of MUFA synthesis and incorporation pathway. SREBP-1c functions as a transcriptional regulator of lipogenic gene expression, including SCD1. SCD1 catalyzes the conversion of saturated fatty acids into MUFAs^28^, while ACSL3 promotes the activation and incorporation of MUFAs into phospholipids^1^. Interestingly, MBOAT1/2 was not affected by LXR activation (Extended Data Fig. 3b). To validate these findings across a broader range of cell models, we analyzed the expression of LXRα, SREBP-1c, SCD1 and ACSL3 also in HT-1080 (Fig. 4b) and HT-29 (Fig 4c), as well as SCD1 and ACSL3 in *ex vivo* primary mouse hepatocytes (Extended Data Fig. 3c). In all cell lines, these genes were upregulated and this induction was markedly attenuated upon co-treatment with the LXR inhibitor (Fig. 4b and 4c). Finally, these findings were validated at the protein level in HepG2, HT-1080, and HT-29 cells. LXR activation increased SREBP-1c, SCD1, and ACSL3 protein expression, an effect that was attenuated by LXRi co-treatment (Fig. 4d, Extended Data Fig. 4a-c). Collectively, these results establish a conserved LXR-MUFA regulatory axis across multiple cell lines.

**Fig. 4.**
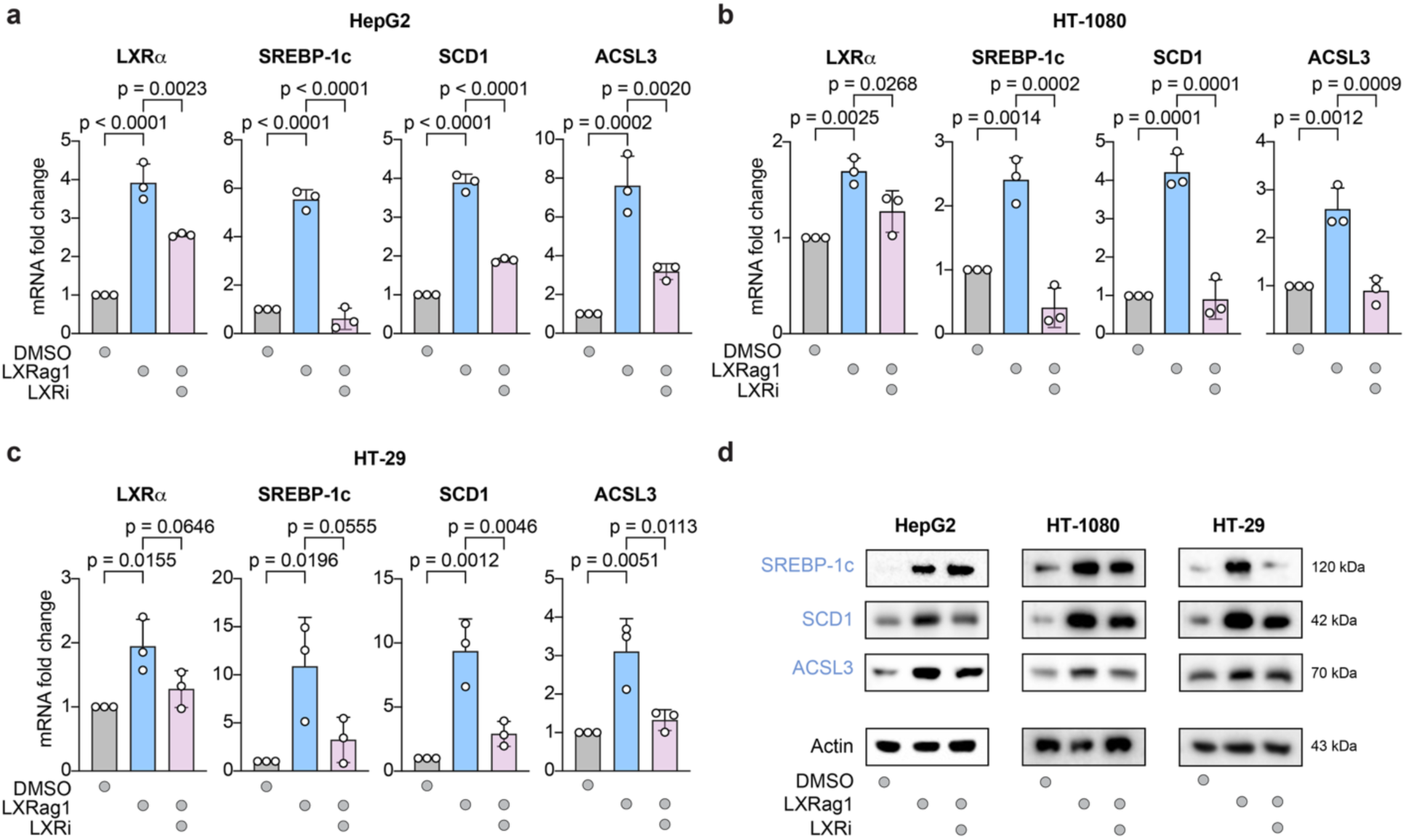
LXR-mediated upregulation of the MUFA-PL synthesis pathway. **a,b,c,** qRT-PCR measurements of LXRα, SREBP-1c, SCD1 and ACSL3 show increased mRNA foldchange upon LXR activation (10 µM LXRag1) and decrease upon co-treatment with LXRi (25 µM) in (a) HepG2, (b) HT-1080 and (c) HT-29 cells; levels of mRNA were normalized to RP2 expression; mean ± SD of *n* = 3 independent biological replicates; ordinary one-way ANOVA with Šidák’s multiple-comparison test. **d,** Western blots show upregulation of SREBP-1c, SCD1, and ACSL3 protein levels upon LXRag1 (10 µM) treatment, which is attenuated by co-treatment with LXRi (25 µM) in HepG2, HT-1080 and HT-29 cells. Protein expression of β-actin was used as control; *n* = 3 independent biological replicates.

### Oxysterol-mediated LXR activation suppresses lipid peroxidation

We subsequently tested whether LXR activation by its endogenous ligand protects cells from ferroptosis. Ferroptotic HepG2 cells were treated with 24(S),25-epoxycholesterol (24,25-EC), a natural oxysterol and ligand of LXR^29^. Consistent with our data using LXRag1, the endogenous oxysterol significantly reduced RSL3-induced lipid peroxidation (Fig. 5a). Importantly, 24,25-EC did not reduce C11-BODIPY oxidation induced by AAPH (Fig. 5b), indicating that its antiferroptotic effect is not due to antioxidant properties. We performed quantitative qRT-PCR in HepG2 cells following 24,25-EC oxysterol treatment to confirm LXR activation and to assess whether endogenous activation regulates the same previously identified target genes (Fig. 4). Treatment with 24,25-EC resulted in mRNA upregulation of LXRα and LXRβ, as well as SREBP-1c, SCD1 and ACSL3 (Fig. 5c), resulting in the upregulation of the MUFA-PL synthesis pathway.

**Fig. 5.**
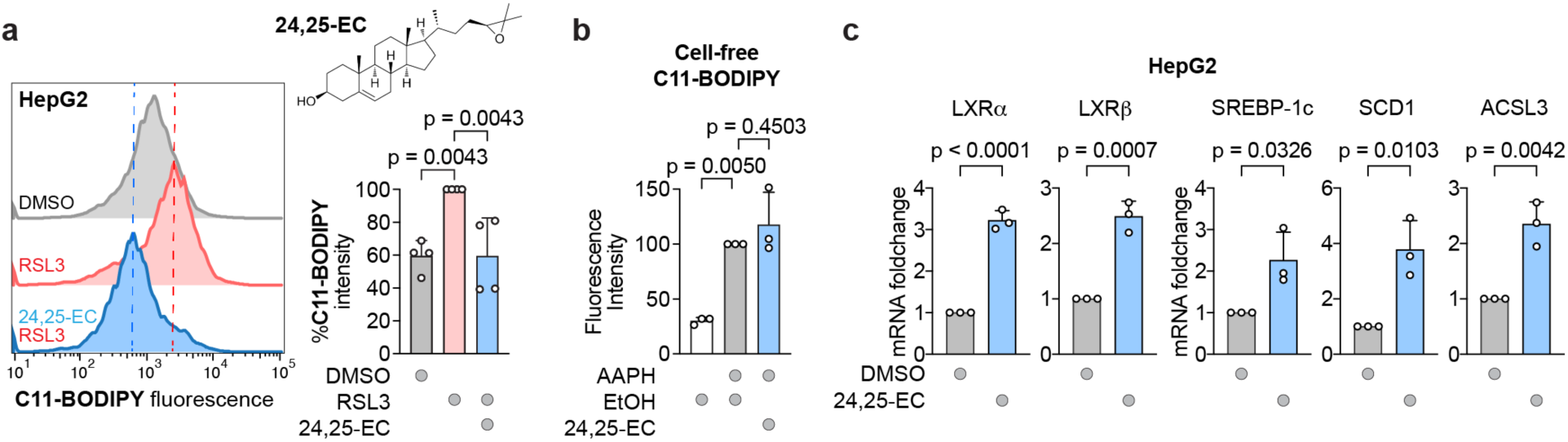
LXR activation by oxysterols reduces lipid peroxidation and upregulates the MUFA-PL synthesis pathway. **a,** Fluorescence shift of C11-BODIPY indicates treatment with 100 nM endogenous LXR-ligand 24(S),25-epoxycholesterol (24,25-EC) decreases lipid peroxidation in ferroptotic (1 µM RSL3) HepG2 cells; mean ± SD of *n* = 3 independent biological replicates; ordinary one-way ANOVA with Šidák’s multiple-comparison test. **b,** Cell-free oxidizable BODIPY-C11 assay treated with 7.5 mM free-radical-producing 2,2′-azobis(2-methyl-propanimidamide) dihydrochloride (AAPH) indicates no antioxidant capacity of 24,25-EC (100 nM); mean ± SD of *n* = 3 independent biological replicates; ordinary one-way ANOVA with Šidák’s multiple-comparison test. **c,** qRT-PCR of LXRα, LXRβ, SREBP-1c, SCD1 and ACSL3 show increased mRNA foldchange upon treatment with 24,25-EC (100 nM); levels of mRNA were normalized to RP2 expression; mean ± SD of *n* = 3 independent biological replicates; unpaired t-test.

### Lipid remodeling to MUFA-rich membranes by LXR counteract ferroptosis

We performed targeted lipidomics to assess how LXR activation by LXRag1 affects cellular lipid composition. Principal component analysis (PCA) revealed a clear separation between DMSO- and LXRag1-treated HepG2 cells (Fig. 6a). Several monounsaturated fatty acids (MUFAs) were increased upon LXR activation, whereas polyunsaturated fatty acids (PUFAs) were reduced (Fig. 6b). Consistently, LXR activation increased the MUFA/saturated fatty acid (SFA) as well as MUFA/PUFA ratios (Fig. 6c), particularly within phospholipid classes phosphatidylethanolamines (PE), phosphatidylcholines (PC), phosphatidylserines (PS), and phosphatidylinositols (PI) (Fig. 6d), which are critical components of cellular membranes and implicated in ferroptosis. In line with this shift, lipidomics-derived SCD1 product-to-precursor ratios showed increased D9D(C16) and D9D(C18) indices following LXR activation, which were attenuated upon co-treatment with the LXR inhibitor (Extended Data Fig. 5a). These changes indicate enhanced synthesis and incorporation of MUFAs, demonstrating a lipid remodeling by LXR activation. The LXRag1-mediated lipid remodeling toward MUFA-enriched membranes suppressed RSL3-induced ferroptosis in HepG2 and HT-1080 cells (Fig. 6e and 6f). However, co-treatment with increasing concentrations of PUFAs counteracted this effect and restored ferroptotic cell death in a dose-dependent manner (Fig. 6e and 6f). ACSL3 catalyzes the final step of MUFA incorporation into membrane phospholipids. To test whether LXR-driven ferroptosis inhibition depends on lipid remodeling, we generated an ACSL3 knockout in HT-1080 cells. While LXRag1 rescued cell viability upon RSL3-induced ferroptosis in control cells, this effect was markedly reduced in ACSL3-KO cells (Fig. 6g). Consistently, the ability of LXRag1 to suppress lipid peroxidation was largely lost upon ACSL3 knockout, indicating that LXR-mediated protection from ferroptosis requires ACSL3-dependent MUFA incorporation (Fig. 6h, Extended Data Fig. 6b-d). To conclude, we show that LXR activation leads to increased expression of SREBP-1c, SCD1 and ACSL3. This promotes lipid remodeling toward MUFA-enriched cell membranes, which are less susceptible to lipid peroxidation and thereby reduce sensitivity to ferroptosis (Fig. 6i).

**Fig. 6.**
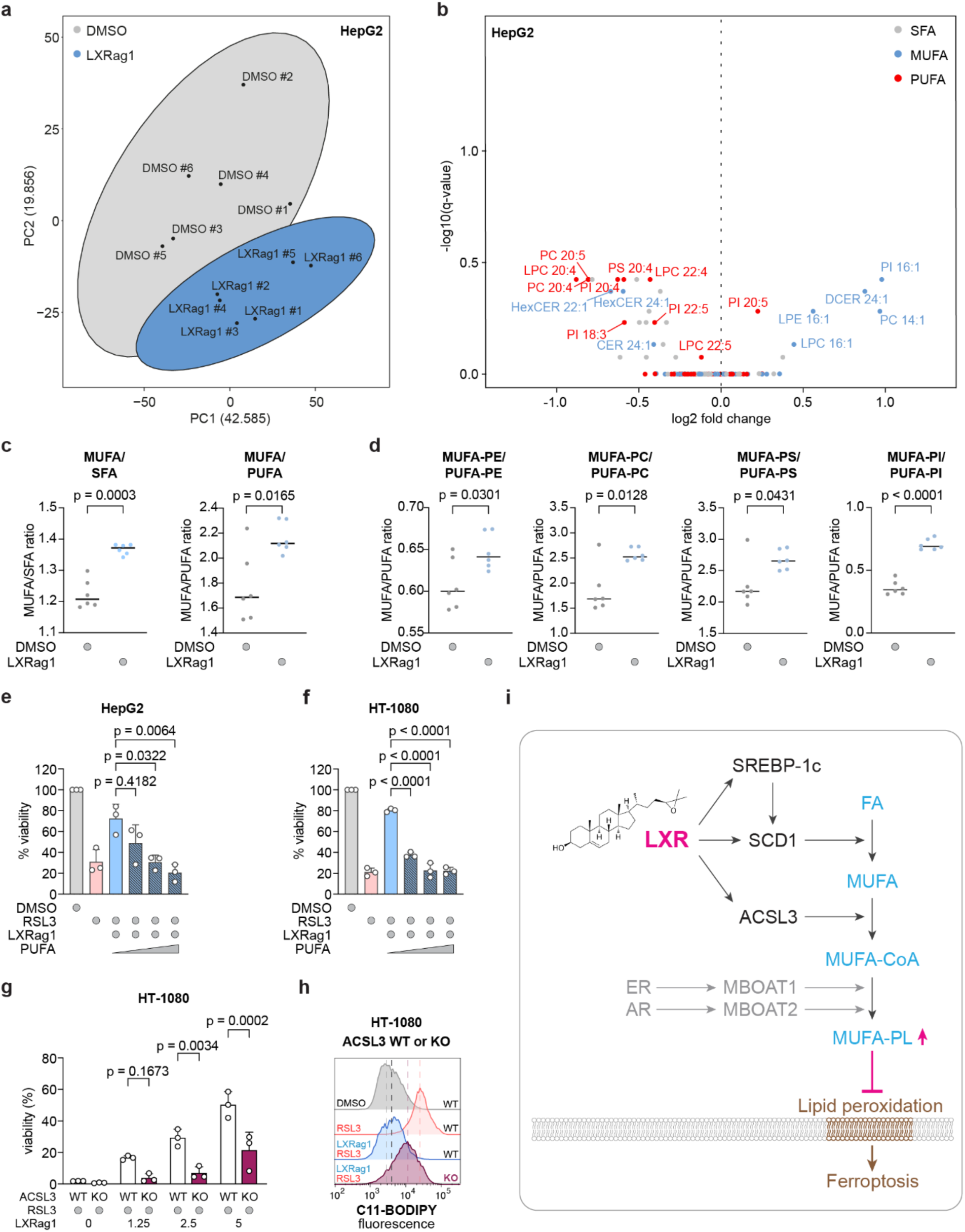
LXR protects against ferroptosis via MUFA-rich membrane remodeling. **a,** Principal component analysis (PCA) of lipidomics raw data comprising six replicates per condition treated with DMSO (control) or LXR agonist 1 (10 µM); data are mean-centered and scaled to unit variance; *n* = 6 independent biological replicates. **b,** Volcano plot showing fold changes upon LXR activation (10 µM LXRag1 vs. DMSO control) reveals increased levels of monounsaturated fatty acids (MUFA); saturated fatty acids (SFA, grey), monounsaturated fatty acids (MUFA, blue) and polyunsaturated fatty acids (PUFA, red); *n* = 6 independent biological replicates. **c,** Fatty acid ratios of MUFA to SFA and MUFA to PUFA display increased MUFA levels in LXRag1 treated HepG2 cells compared to control cells; mean ± SD of *n* = 6 independent biological replicates, Welch’s t-test. **d**, Increased MUFA/PUFA ratio observed in lipid classes phosphatidylethanolamine (PE), phosphatidylcholine (PC), phosphatidylserine (PS), phosphatidylinositol (PI); mean ± SD of *n* = 6 independent biological replicates, Welch’s t-test. **e**,**f**, Increase in viability observed in LXRag1-treated ferroptotic (150 nM RSL3) cells is progressively reduced with increasing concentrations of PUFA (1.5 µM, 6 µM, 12.5 µM) in (e) HepG2 and (f) HT-1080 cells. **g**, Dose-dependent increase in viability in LXRag1-treated (1.25 µM, 2.5 µM, 5 µM) wildtype HT-1080 cells exposed to 150 nM RSL3 is attenuated in ACSL3-KO HT-1080 cells treated under the same conditions; mean ± SD of *n* = 3 independent biological replicates; ordinary one-way ANOVA with Tukey’s multiple-comparison test. **h**, Fluorescence shift of C11-BODIPY in LXRag1 treated HT-1080 wildtype cells towards a lowered lipid peroxidation is attenuated in ACSL3-KO cells, *n* = 3 independent biological replicates. **i,** Graphical illustration of oxysterol-LXR mediated ferroptosis inhibition.

Thus, oxysterol-activated LXR couples ligand sensing to a selective lipid-remodeling program rather than broadly engaging ferroptosis defense systems.

## Discussion

In this study, we identified LXR as a novel oxysterol-sensing suppressor of ferroptosis through a chemical genetics screen. Mechanistically, oxysterol-activated LXR promotes a MUFA-enriched membrane composition through selective induction of SREBP-1c, SCD1 and ACSL3, thereby reducing susceptibility to lipid peroxidation. Reintroduction of PUFAs following LXR-mediated lipid remodeling reduces the protective effect against ferroptosis, and ACSL3 knockout attenuates LXR-mediated ferroptosis resistance, directly linking LXR-mediated lipid remodeling to ferroptosis protection. Conversely, LXR inhibition or knockout sensitized cells to ferroptotic cell death, providing a potential means of targeting LXR-dependent cancers. This pathway is robust across multiple model systems, including cancer cell lines, 3D spheroids, and *ex vivo* hepatocytes. Previous studies established LXR as a regulator of fatty acid desaturation and phospholipid homeostasis through SCD1- and ACSL3-dependent programs in macrophages and placental trophoblast cells^30, 31^, and our data extend this function to ferroptosis regulation. While oxysterol-mediated sensitization to ferroptosis independent of LXR signaling has been reported^32^, our findings identify endogenous LXR activation as a protective pathway suppressing ferroptosis through adaptive MUFA remodeling. Of note, chronic exposure to 27-hydroxycholesterol was shown to inhibit ferroptosis by stabilizing GPX4 expression^33^, a mechanism distinct from the lipid remodeling program described here.

Recently, we showed that the farnesoid X receptor (FXR), another sterol-related nuclear receptor, suppresses ferroptosis by upregulating a broad panel of ferroptosis-inhibitory genes, including GPX4, FSP1, PPARα, SCD1, and ACSL3^19^. FXR therefore engages several anti-ferroptotic defense systems in parallel. LXR acts through a different principle: first, the activating signals are distinct, as FXR is a sensor for bile acids, whereas LXR is a sensor for oxysterols, some of which arise from radical- and ROS-mediated oxidation of cholesterol^34^; second, the transcriptional output is specific, as LXR activation selectively induces SREBP-1c, SCD1, and ACSL3, while GPX4, FSP1, and MBOAT1/2 are not regulated in an LXR-dependent manner (Fig. 4, Extended Data Fig. 3). Hence, FXR and LXR represent mechanistically distinct nuclear receptors. As ligand-activated transcription factors, nuclear receptors function as cellular sensors of endogenous metabolites to maintain metabolic homeostasis and adapt to metabolic stress^35^. In this context, nuclear receptors may integrate metabolic cues to regulate and fine-tune lipid peroxidation and ferroptosis^1^. LXR may sense oxidative lipid stress and induce a transcriptional program that limits oxidative membrane damage and ferroptotic cell death under conditions of increased lipid peroxidation. Together, these findings identify LXR as a mechanistically distinct nuclear receptor that expands the concept of metabolite-driven ferroptosis regulation.

Dysregulation of these sensing pathways contributes to metabolic disease development and progression. Non-alcoholic fatty liver disease (NAFLD) represents a spectrum of conditions driven by hepatic steatosis, which can progress to the more severe form non-alcoholic steatohepatitis (NASH), associated with advanced liver pathologies including cirrhosis and hepatocellular carcinoma^36^. Ferroptosis has emerged as a potential driver of this transition^37^, and inhibition of ferroptosis ameliorates hepatic steatosis and inflammation in NASH mouse models ^38^. In parallel, oxysterols have been implicated in modulating hepatic disease progression, with reduced oxysterol levels exacerbating hepatic steatosis under obesogenic conditions^39^. Given these connections, LXR activation by oxysterols may provide an additional hepatoprotective mechanism by limiting ferroptotic cell death, while reduced LXR signaling may conversely contribute to NASH progression. In addition, the negative correlation between LXRα expression and patient survival in HCC is consistent with our mechanistic findings: high LXR activity suppresses ferroptosis, providing tumor cells with a survival advantage that promotes tumor progression and worsens prognosis. This highlights LXR inhibition as a potential therapeutic strategy to restore ferroptosis sensitivity in LXRα-high tumors, as supported by our finding that LXRi sensitizes ferroptosis-resistant cells to RSL3-induced cell death. Hence, future studies in relevant *in vivo* models will address these hypotheses and evaluate the therapeutic potential of pharmacological LXR modulation in ferroptosis-associated liver diseases including NASH and cancer.

Together, our findings identify the liver X receptor as a nuclear receptor that couples oxysterol signaling to adaptive membrane remodeling and ferroptosis resistance.

## Methods

### Cell culture

The following immortalized cell lines were used for this study: human fibrosarcoma (HT-1080), human hepatocellular carcinoma (HepG2), colorectal adenocarcinoma (HT-29) and mouse embryonic fibroblasts (MEF). HT-1080, HepG2 and HT-29 were purchased from ATCC, MEF were a gift from Prof. Dr. Krappmann (Helmholtz Munich).

All cell lines were cultured in Dulbecco’s Modified Eagle’s Medium (DMEM, ThermoFisher Scientific, 41966-029) supplemented with 10% fetal bovine serum (FBS, ThermoFisher Scientific), 1% Penicillin-Streptomycin (ThermoFisher Scientific) and 1% non-essential amino acids (NEAA, ThermoFisher Scientific) in a humidified incubator at 37°C and 5% CO_2._ To minimize clumping, HepG2 cells were strained through a 70 µM cell strainer prior to seeding. Cells were regularly tested (including mycoplasma) and were completely contamination-free.

### Compounds

The following compounds were purchased: (1S,3R)-RSL3 (Sigma), imidazole ketone erastin (IKE, Cayman Chemical), ML210 (Sigma), Ferrostatin-1 (Fer-1, Sigma), GSK3987 (LXRag1, Biotrend), AZ876 (Biotrend), GSK2033 (LXRi, Biotrend), HX531 (Biotrend), 24(S),25-epoxycholesterol (24,25-EC, Abcam), Staurosporine (Stauro, TargetMol), z-VAD-FMK (zVAD, TargetMol), LCL161 (MedChemExpress), tumor necrosis factor alpha (TNFα, biomol), Necrostatin-1 (Nec-1, BioVision). All compounds except 24(S),25-epoxycholesterol were dissolved in dimethyl sulfoxide (DMSO). 24(S),25-epoxycholesterol was dissolved in 100% ethanol.

### Chemical-genetic screening

For screening, HT-1080 cells were seeded in white 384-well culture plates (CulturPlate, 6007680, Revvity) at a density of 750 cells per well. After 24h of incubation, a library of 550 activators and inhibitors of nuclear receptors (HY-L126, MedChemExpress; in DMSO at a stock concentration of 1 mM) was applied to the cells using a Sciclone G3 Liquid Handler (PerkinElmer). A total volume of 0.5 µL compounds were used to achieve a final concentration of 10 µM. Subsequently, ferroptosis was induced by treating cells with 100 nM RSL3. As controls, cells were either treated with DMSO only or treated with 2 µM Ferrostatin-1 (Fer-1). After 18 hours of ferroptosis induction, cell viability was assessed by adding 20 µL of CellTiter Glo 2.0 (G9243, Promega) and measuring luminescence in an EnVision 2104 Multilabel plate reader (PerkinElmer). Hits were defined as a luminescence signal higher than a threshold of three times the standard deviation from the median of the compound-treated population.

### Kaplan-Meier Plotter

Liver hepatocellular carcinoma was analyzed for pan-cancer survival analysis using the KM Plotter mRNA (RNA-seq) module of the Kaplan-Meier Plotter (kmplot.com)^25–27^ with data accessed in May 2026. Patients were stratified based on NR1H3 (LXRα) expression levels using a median split cut-off. Kaplan-Meier survival curves were generated to evaluate the association between NR1H3 expression and overall survival (*n*=371). Statistical significance was assessed using the log-rank test, and hazard ratios (HR) with corresponding 95% confidence intervals were obtained to estimate the prognostic impact of NR1H3.

### Viability Assays

To assess viability, cells were seeded in white 384-well culture plates (CulturPlate, 6007680, Revvity) at a density of 750 cells (HT-1080 and HT-29) or 1000 cells (HepG2) per well. After 24 hours of incubation, cells were treated with 12-point serial dilutions of ferroptosis inducers or inhibitors with indicated concentrations. DMSO was used as a control treatment for normalization of luminescence signals. After a treatment duration of 18 hours, viability was measured using CellTiter Glo 2.0 (G9243, Promega) according to the manufacturer’s instructions and reading out luminescence signal in a EnVision 2104 Multilabel plate reader (PerkinElmer). To assess apoptotic cell death, HT-1080 were seeded in a similar fashion and treated with 1 µM staurosporine diluted in culture medium before adding 50 µM z-VAD-FMK as apoptosis inhibitor or LXR agonists. After 24 hours, activity of caspases 3 and 7 was measured with the Caspase-Glo® 3/7 Assay Reagent (Promega) according to the manufacturer’s instructions. To assess necroptotic cell death, MEF cells were seeded in a similar fashion and treated with a cocktail of 20 ng/mL TNFα, 10 µM LCL161 and 10 µM z-VAD-FMK diluted in culture medium to induce necroptosis before adding necrostatin-1 as an inhibitor or LXR agonists. Viability was measured after 18 hours by adding CellTiter Glo 2.0 according to the manufacturer’s instructions and reading out luminescence in an EnVision 2104 Multilabel plate reader (PerkinElmer).

### Ex vivo mouse experiments

Livers from *Hfe* mice (34-65 weeks old) were perfused through the vena cava in a single-pass manner using calcium-free buffer (HBSS without calcium and magnesium, supplemented with 0,9 mM MgCl₂, 0.5 mM EGTA, and 25 mM HEPES, pH 7.4, Sigma-Aldrich), followed by digestion buffer (HBSS with calcium and magnesium, 25 mM HEPES) and a collagenase solution (190 U/mL collagenase in digestion buffer). All buffers were maintained at 37 °C. After perfusion, livers were excised and transferred to ice-cold isolation medium (Leibovitz L15 (Gibco™, ThermoFisher Scientific), 1% PenStrep (Life Technologies)). Hepatocytes were released by gentle agitation, filtered through a 70 µm nylon mesh, and collected by centrifugation (50 x g, 5 min, 4 °C). Cells were washed with Hepatocyte Wash Medium (Gibco™, ThermoFisher Scientific) with 1% PenStrep and purified on a 90% Percoll monolayer. Viability and yield were assessed prior to seeding in Williams E medium containing 5% FCS, 1% PenStrep, 100 nM dexamethasone (Sigma-Aldrich), and 100 nM insulin (Sigma-Aldrich). After 4 h, the medium was replaced with serum-free Williams E medium (Gibco™, ThermoFisher Scientific) containing 1% PenStrep, 100 nM dexamethasone, and 100 nM insulin. Cells were seeded at a density of 40,000 cells per well in 96-well plates for viability assays and 800,000 cells per well in 6-well plates for qRT-PCR experiments. Cells were treated with 10 µM GSK3987 and with or without 25 µM GSK2033 for 18 hours. DMSO-only treatment served as a control.

### Spheroid imaging

HT-1080 cells were seeded into a 96-well round bottom ultra-low attachment microplate (7007, Corning costar) at a density of 2000 cells per well. After 48 hours of incubation at 37°C cells were treated with 200 nM RSL3, 10 µM GSK3987, 2 µM Ferrostatin-1 or DMSO as control. Nuclei were stained with Hoechst 33342 (Sigma Aldrich) at a dilution of 1:10,000 after 48 hours, followed by 30 min incubation at 37°C. Spheroids were imaged using the operetta high content screening system (PerkinElmer) at 10x magnification. Image analysis was performed using the Harmony software (PerkinElmer).

### Crystal violet staining

HT-1080 cells were seeded into a 6-well plate at a density of 200,000 cells per well and treated with 150 nM RSL3, 10 µM GSK3987 and with or without 25 µM GSK2033 for 18 hours. DMSO was used as a control treatment. Two hours later, the medium was removed and cells were washed with Milli-Q water. 1.5 mL of crystal violet staining solution (0.5% crystal violet in 20% methanol) was added per well. Cells were incubated at room temperature for 20 min and rinsed with Milli-Q water until no excess stain remained. Images were quantified using ImageJ.

### Transfection overexpression plasmids

LXRα (NM_005693) and LXRβ (NM_007121) expression plasmids were purchased from OriGene. HT-1080 cells were seeded in 6-well plates at a density of 200,000 cells per well. For transfection, a total of 1 µg plasmid DNA per well (0,5 µg LXRα plasmid and 0,5 µg LXRβ plasmid) was diluted in 90 µL Opti-MEM reduced-serum medium (31985062, ThermoFisher Scientific). Subsequently, 3 µL X-tremeGENE HP DNA transfection reagent (6366244001, Merck) was added, and the mixture was incubated for 15 min at room temperature. The transfection mixture was added dropwise to the cells, which were incubated for 24 hours under cell culture conditions.

### Electroporation of siRNA

Electroporation of siRNA was performed using the Neon NxT Electroporation System (ThermoFisher Scientific). HT-1080 LXRα-knockout cells were cultivated and harvested using 0.25% Trypsin-EDTA (25200056, ThermoFisher Scientific) for 2 min. The desired number of cells was transferred to a falcon tube and centrifuged for 5 min at 400 × g at room temperature. Cells were washed with PBS without Ca²⁺ and Mg²⁺ and centrifuged again for 5 min at 400 × g at 4 °C. Cell pellets were kept on ice until electroporation. Post-electroporation culture vessels were prepared by filling them with pre-warmed antibiotic-free culture medium and pre-incubating them at 37 °C and 5% CO₂. The Neon NxT pipette station was prepared according to the manufacturer’s instructions. Cell pellets were resuspended in R buffer at a final density of 3 × 10⁵ cells per 10 µL, and 100 nM LXRβ siRNA (sc-45316, Santa Cruz Biotechnology) or negative control siRNA (SR-CL000-005, Eurogentec) was added and gently mixed. A total volume of 100 µL was aspirated into a Neon NxT Tip, and electroporation was performed using a single pulse with a pulse voltage of 950 V and a pulse width of 50 ms. Electroporated cells were transferred to the prepared culture vessels and incubated for 48 h under cell culture conditions.

### Generation of KO cell lines

Lentiviral vectors were generated using a third-generation lentiviral packaging system. HEK-293T cells were transfected with X-tremeGENE HP DNA Transfection Reagent (6366546001, Sigma-Aldrich) together with a plasmid mixture consisting of pLentiCRISPR v2 carrying sgRNAs targeting human ACSL3 (sg1: CACCGTGGTGAAGAGTAACCAATG; sg2: CACCGGGCTGGAACAATTTCCGA; sg3: CACCGGGGTGAAATTCTTATTGG), pMDLg/pRRE (12251, Addgene), pRSV-Rev (12253, Addgene), and pCMV-VSV-G (8454, Addgene). Lentiviral-containing supernatants were collected 48 hours after transfection and filtered through a 0.45 μm filter. HT-1080 cells were transduced with the filtered supernatants (8 μg/mL protamine sulfate). The medium was replaced with fresh medium supplemented with blasticidin (10 μg/mL) 48 hours post-transduction. Cells were maintained in antibiotic selection for one week. All lentiviral production and transduction procedures were conducted under biosafety level 2 (BSL-2) conditions.

### qRT-PCR

The total RNA was isolated using the Monarch Total RNA Miniprep Kit (T2110S, New England BioLabs) according to the manufacturer’s protocol. Genomic DNA was removed using the Monarch SC2 Spin Columns (T3017L, New England BioLabs). For cDNA synthesis, oligo(dT)₁₈ primers (SO132, ThermoFisher Scientific) and random hexamer primers (SO142, ThermoFisher Scientific) were used together with the Maxima H Minus Reverse Transcriptase (EP0752, ThermoFisher Scientific). Reverse transcription was performed with an optimized incubation of 10 min at 21°C followed by 30 min at 50°C. RT-qPCR was carried out using PowerUp SYBR Green Master Mix (ThermoFisher Scientific) on a LightCycler 480 system (Roche). Gene expression levels were normalized to RNA polymerase II (RP2) expression and quantified using the ΔΔCt method. A list of primers used for qRT-PCR is provided in the Supplementary Table 3.

### Western Blot

HepG2, HT-1080 or HT-29 cells were plated in 6-well plates at a density of 200,000 cells per well and after a 24-hour incubation treated with 10 µM GSK3987, 25 µM GSK2033 or DMSO as a vehicle control for 18 hours. Cells were subsequently collected in 2x reducing Roti-Load buffer (K929.1, Carl Roth) using a cell scraper and lysed by sonication with an ultrasonic processor (Hielscher UP200S). Proteins were denatured by heating the samples at 95 °C for 5 minutes. Protein extracts were separated on NuPAGE™ 4–12% Bis-Tris gels (NP0329BOX, Invitrogen) using 1x MOPS SDS running buffer (NP0001, Invitrogen) and transferred onto a PVDF membrane via semi-dry blotting. Membranes were blocked for 30 min at room temperature in TBS-T containing 5% milk powder (T145.3, Carl Roth) and then incubated overnight at 4 °C with primary antibodies (mouse anti-β-Actin antibody (8H10D10), 12262S, Cell Signaling Technology; mouse anti-β-actin antibody (ACTB), A5441, Sigma-Aldrich; rabbit anti-ACSL3, ab151959, abcam; rabbit anti-ACSL3 antibody (E2S9L), 83319T, Cell Signaling Technology; rabbit anti-SCD1 antibody (EPR21963), ab236868, abcam; mouse anti-SREPB1 (2A4), MA5-11685, Invitrogen; rabbit anti-Glutathione Peroxidase 4 antibody (EPNCIR144), ab125066, abcam; mouse anti-GCH1 antibody (OTI5A1), ab236387, abcam; rat anti-AIFM2 (6D8-1-1), Helmholtz Munich antibody core facility; rabbit anti-LXRα, PA1-330, Invitrogen; goat anti-LXRα + LXRβ antibody, ab24362, abcam). Afterwards, the membrane was washed three times for 10 minutes in TBS-T and incubated for one hour at room temperature in secondary antibody (anti-Mouse IgG (H+L), HRP Conjugate, W4021, Promega, anti-Rabbit IgG (H+L), HRP Conjugate, W4011, Promega, goat anti-Rat IgG (H+L), AB_2338128, Jackson ImmunoResearch; donkey anti-Goat IgG (H+L), AB_2313587, Jackson ImmunoResearch) diluted 1:2,500 in 5% milk-TBS-T. After washing three times for 10 min in TBS-T, chemiluminescent signals were detected using Western Lightning ECL Pro (NEL120001EA, Revvity) Sapphire Biomolecular Imager (Biozym). Full-scan western blots are provided in Extended Data Fig. 6-8.

### Flow cytometry

HepG2, HT-1080, and HT-29 cells were seeded in 6-well plates at a density of 200,000 cells per well and incubated for 24 hours. Cells were treated with RSL3 for 2 hours (HT-1080, HT-29) or 3 hours (HepG2) in the presence of the indicated compounds. Lipid peroxidation was assessed by either BODIPY 581/591 C11 staining or 4-hydroxynonenal (4-HNE) immunostaining. For BODIPY staining, cells were incubated with BODIPY 581/591 C11 (D3861, Invitrogen; 2 µM) for 30 minutes at 37 °C prior to harvest. Cells were harvested using 0.25% Trypsin-EDTA (25200056, ThermoFisher Scientific) and washed twice with PBS. For immunostaining cells were blocked in 10% normal goat serum (50197Z, ThermoFisher Scientific) for 30 minutes on ice and incubated with an anti-4-HNE antibody (ab46545, Abcam; 1:50 in 1% BSA in PBS) for one hour on ice, followed by an anti-rabbit Alexa Fluor 488 secondary antibody (A32731, ThermoFisher Scientific; 1:200 in 1% BSA in PBS) for 30 minutes on ice. After washing twice with PBS cells were resuspended in 300 µL PBS and analyzed by flow cytometry. For PI staining, cells were harvested as described, washed with PBS, and resuspended in PBS containing 1 µg/mL PI. A total of 10,000 events per condition were recorded using an Attune acoustic flow cytometer (Applied Biosystems) in the BL-1 channel for C11-BODIPY and 4-HNE or BL-3 channel for PI. Data were analyzed using FlowJo v10.8.1 (BD Life Sciences).

### Cell-free BODIPY assay

GSK3987 and 24(S),25-epoxycholesterol were prepared in 150 µL PBS at a final concentration of 25 µM. For vehicle control, an equivalent volume of DMSO or EtOH were added to 150 µL PBS. BODIPY 581/591 C11 (Thermo Fisher Scientific) was diluted in PBS to a final concentration of 1.875 µM, while 2,2’-Azobis(2-methylpropionamidine) dihydrochloride (AAPH, Sigma) was prepared at 7.5 mM in 150 µL PBS. A non-oxidized control containing DMSO or EtOH, but no AAPH was included. The respective solutions were combined, loaded into a black 96-well plate (Greiner Bio-One) and fluorescence was measured every 5 min for a total duration of 2 hours at 495 nm/520 nm using a PerkinElmer EnVision 2104 Multilabel plate reader.

### Lipidomics

Cells were seeded in 10-cm culture dishes at a density of 1 Mio cells per dish. After 24 hours, cells were treated for 18 hours with either vehicle control (DMSO) or the LXR agonist GSK3987 (10 µM), with six independent biological replicates per condition. To minimize metabolic activity, cells were harvested on dry ice. Culture dishes were washed twice with 11 mL ice-cold PBS (dry ice), and residual PBS was completely removed. Subsequently, 800 µL of ice-cold 80% methanol (HPLC grade), pre-cooled overnight at −80 °C in a glass flask, was added directly to the cells and evenly distributed. Cells were scraped using a rubber-tipped scraper, and the extracts were transferred into collection tubes kept on dry ice. Remaining material was recovered with an additional 200 µL of 80% methanol and combined with the initial extract. Samples were stored at −80 °C until lipidomics analysis. Cell samples were homogenized in 80% MeOH with 320 mg glass beads (0.5 mm, VK-05, PeqLab) using a PeqLab Precellys24 homogenizer. Samples were cooled to 0–3 °C and homogenized twice at 5500 rpm for 25 s with 5 s breaks in-between. The homogenate was further used for two purposes. First, the cell count was estimated using DNA levels based on fluorescence labeling with the Hoechst dye (final concentration: 20 µg/mL in PBS) as previously described^40^. Second, samples were extracted for lipidomics analyses as described in the following paragraph.

The lipid extraction is based on the protocol by Matyash *et al.*^41^. After thawing at room temperature (RT) for 30 min, 200 µL of the cells samples were transferred into 1.5 mL glass vials together with 60 µL of MilliQ water (H2O). For accurate quantification, 25 µL of a mix of 53 deuterated internal standards were added to the samples (Ultimate SplashOne, dFA 18:1, dCer d18:0/13:0, Glu Cer(d18:1-d7/15:0), dLacCer d18:1/15:0, 15:0-18:1-d7-PA. For lipid extraction 575 µL methyl *tert*-buthyl ether (MTBE, LC grade) were added followed by incubation for 30 min on an orbital shaker DOS-10L (Neolabline, Heidelberg, Germany) at 300 rpm. For phase separation, 200 μL of MS-grade H_2_O was added to each vial. The mixtures were vortexed, and the vials were centrifuged at 5,000 x g for 10 min at RT with a Sigma 4-5C centrifuge (Qiagen, Hilden, Germany). The upper (organic) phase was transferred into new glass vials and evaporated with nitrogen gas using a TurboVap® 96 dual evaporator (Biotage, Uppsala, Sweden). The aqueous phase was again extracted with 100 µL MeOH and 300 µL MTBE. After addition of 100 µL H_2_O, the samples were incubated for 5 min at RT at 300 rpm and then centrifuged for 10 min at 5,000 x g. The organic phase was transferred into the respective vial from the first extraction step and evaporated to dryness with gaseous nitrogen. Samples were reconstituted in 825 µL running solvent (10 mM ammonium acetate in Dichloromethane:MeOH (50:50, v/v)). For quality control purposes, 60 µL of each study sample were pooled in a 1.5 mL Eppendorf tube (QC-pool samples). After vortexing, 200 µL aliquots were created and extracted with the above-described procedure. Additionally, three blank samples consisting of 200 µL 80% MeOH and three aliquots à 25 µl of a commercial pooled human plasma sample were prepared and extracted as described above, with the exception that 75 µL of H_2_O were added to the commercial plasma samples during the first extraction step.

Lipidomic analyses of the cell samples were performed using the Differential Ion Mobility Shotgun Lipidomics Assay (DMS-SLA)^42^. An extensive description of the experimental details has previously been published^43^. All samples were measured with a SCIEX Exion UHPLC-system coupled to a SCIEX QTRAP 6500+ mass spectrometer equipped with a SelexION differential ion mobility interface (SCIEX, Darmstadt, Germany) operated with Analyst 1.6.3. Per run, 225 µL of the re-dissolved sample were injected using the running solvent (10 mM ammonium acetate in Dichloromethane:MeOH (50:50, v/v)) at an isocratic flow rate of 24 µL/min. After 9 minutes the flowrate was ramped to 90 µL/min for 2 min to allow for washing. Each sample was analyzed using multiple reaction monitoring (MRM) in two consecutive flow injection analysis (FIA) runs. In the first run, phosphatidylcholines (PC), phosphatidylethanolamines (PE), phosphatidylglycerols 2 (PG), phosphatidylinositols (PI), phosphatidylserines (PS), and sphingomyelins (SM) were separated with the SelexION DMS cell using field asymmetric ion mobility mass spectrometry (FAIMS) prior to analysis in the Turbo Spray IonDrive source of the mass spectrometer. To enhance the separation of the lipid classes, 1-propanol was used as a chemical modifier. In the second run, cholesteryl esters (CE), ceramides (Cer d18:1), dihydroceramides (Cer d18:0), lactosylceramides (LacCER), hexosylceramides (HexCER), phosphatidic acid (PA), lysophosphatidylcholines (LPC), lysophosphatidylethanolamines (LPE), lysophosphatidylglycerols (LPG), lysophosphatidylinositols (LPI), lysophosphatidylserines (LPS), free fatty acids (FA), diacylglycerols (DG), and triacylglycerols (TG), were measured with the DMS-cell switched off. The mass spectrometer was operated with the following conditions: curtain gas 20 psi, ion source gas 1 14 psi, ion source gas 2 20 psi, Collision gas medium, temperature 150 °C, separation voltage 3500 V, ion spray voltage +4200 and +4500 V in ESI+ mode and − 4400 and −3300 V in ESI− mode for run 01 and 02, respectively. Prior to each batch, the DMS-cell was tuned, and the stability and sensitivity of the instrument was checked with the EquiSPLASH mixture (AvantiPolar) by using the Shotgun Lipidomics AssistantSLA software (SLA.v1.5; github.com/syjgino/SLA/tree/v1.5-keyV4). Sciex wiff files were converted to mzml format using the Proteowizard msconvertGUI tool (v3.0.22074; proteowizard.sourceforge.io/dow nload.html). The converted files were subsequently processed using the SLA software (SLA.v1.5; github.com/syjgino/SLA/tree/v1.5-keyV4). The average peak intensity was calculated from 20 scans, as per MRM. An MRM was excluded if either an analyte or its corresponding internal standard showed zero intensity in more than 2 out of the 20 scans. Lipid species concentrations were calculated from the ratio of the averaged intensity of an analyte to its internal standard of known concentration using standard_dict-V4_1.3. Lipid species concentrations were subsequently corrected for Type-II isotopic overlap using lipid specific correction factors as listed in ISOcorrectlistV4_1.3_20220926 and are reported in nmol/ml homogenate. The shotgun lipidomics raw data set contained 1202 individual lipid species. Data was subsequently pre-processed using R (version 4.4.1). To assure high data quality, a multi-step procedure was applied: In the first step of this quality control (QC) procedure, lipids with missing values in more than 35% in the pool samples were discarded from the data set (n = 173). In the second step, the group-specific missingness was evaluated i.e., whether a specific lipid is observed in only one of the biological groups. Lipids exhibiting a groupwise missingness of 50% in all groups were discarded from the data set (n = 2). In the following filtering step, ether-linked PE species (PE-O) having an isobaric interference of > 10 % with the respective odd chain PE were marked as “contaminated”. In case a PE-O species was contaminated in > 20 % of the samples per group in all groups under investigation, the PE-O species was removed from the data set (n = 8). We next evaluated the signal-to-noise ratio (S/N) which was calculated as the ratio between the analyte concentration in the sample matrix and the mean concentration of the same analyte in the blank samples. A lipid was removed in case S/N > 3 in more than 50 % of the samples per group (n = 127). Next, lipids with a coefficient of variation (CV) > 25 %, determined by the QC-pool samples, were removed from the data set (n = 22). The last quality control step comprised the calculation of the dispersion ratio (D-ratio) that uses the ratio of the technical variance (determined by QC-pool samples) and the total variance (variance of the biological samples) as a quality marker for each lipid^44^. We used a D-ratio cut-off of 50%, as this implies that the technical variance is larger than the biological variance (n = 85 lipids were removed). After quality control, 785 lipid species remained in the cells data set, which contained 213 missing values (equivalent to 1.1 % of the total data set). Missing values were imputed using the GSimp imputation approach^45^. The algorithm was initialized using a quantile regression approach for left-censored missing data (QRILC). Missing values were then predicted with the Gibbs sampler approach using an elastic net model with alpha = 0.1 and lambda = 0.01. Iterations were used to optimize the value for each missing variable (iters_each) and 10 iterations for the whole data set (iters_all). The classification and abbreviation system for lipid nomenclature follows the LIPID MAPS guidelines^46^. In this system, a lipid class (such as PC, PE, CE, etc.) is indicated first, followed by the associated fatty acid(s) (FA). The fatty acids are denoted by the number of carbon atoms (XX) and double bonds (Y), separated by a colon (for example, CE 16:0). In lipids containing two FA, an underscore (_) is used to separate them when the sn-position is unknown, whereas a slash (/) is used when the sn-position is known. Additionally, ether lipids include an O-(plasmanyl) or P-(plasmenyl) tag, as seen in examples like PE P-18:1/16:1. Triacylglycerols can only be identified with the total number of carbon atoms and double bonds as well as one FA (e.g. TG 52:3-FA16:0). A table with Goslin2.0^47^ normalized lipid names is attached (Supplementary Table 4). Chemicals used were from Avanti Polar Lipids: Ultimate SplashOne (#330820), dFA 18:1 (#861809), dCer d18:0/13:0 (#330726), dGluCer(d18:1-d7/15:0) (#330729), dLacCer d18:1/15:0 (#330727), 15:0-18:1-d7-PA (#791642), EquiSPLASH (#330731).

## Statistical Analysis

Statistical analyses were performed using the GraphPad Prism Software (version 10.6.1). All relevant statistical information is provided in the respective figure legends.

## Data availability

All data are available in the article, its Supplementary Information, and the online source data file. Full scan images of Western blots are provided in Extended Data Figs. 6-8. The compound Library used in Fig. 1b can be found in Supplementary Table 1. Lipidomics results shown in Fig. 6a–d is provided in Supplementary Table 2. Source data are provided with this paper.

## Acknowledgements

We thank Stefanie Brandner and Tamara Rieder for excellent technical assistance.

This work is in part funded by the Deutsche Forschungsgemeinschaft (DFG, German Research Foundation) – TRR 387/1 – 514894665 to KH, JPFA and MD.

## Author contributions

K.H. initiated the study; K.H. oversaw and supervised the whole study; A.K., J.T., S.S., J.S., F.R., O.H., H.H., E.E. performed experiments. A.K., J.T., S.S., O.H., I.R., M.H. analyzed data; K.H., H.Z., M.D., J.P.F.A. supervised the research. A.K., K.H. wrote the initial paper draft; all authors read, commented, and approved the manuscript.

## Responsible use of Artificial Intelligence

Artificial intelligence (AI) tools including ChatGPT, Gemini and DeepL were employed in this manuscript to refine grammar, spelling, and readability. These tools were used exclusively for linguistic enhancement without altering the scientific content, interpretation, or conclusions. All intellectual contributions, data analyses, and critical discussions remain the work of the authors. The final manuscript was thoroughly reviewed and approved by all co-authors to ensure accuracy and integrity.

