## Supplementary Figures for "Oxysterol-sensing by Liver X receptor counteracts ferroptosis via lipid remodeling"

#### Supplementary Figure 1

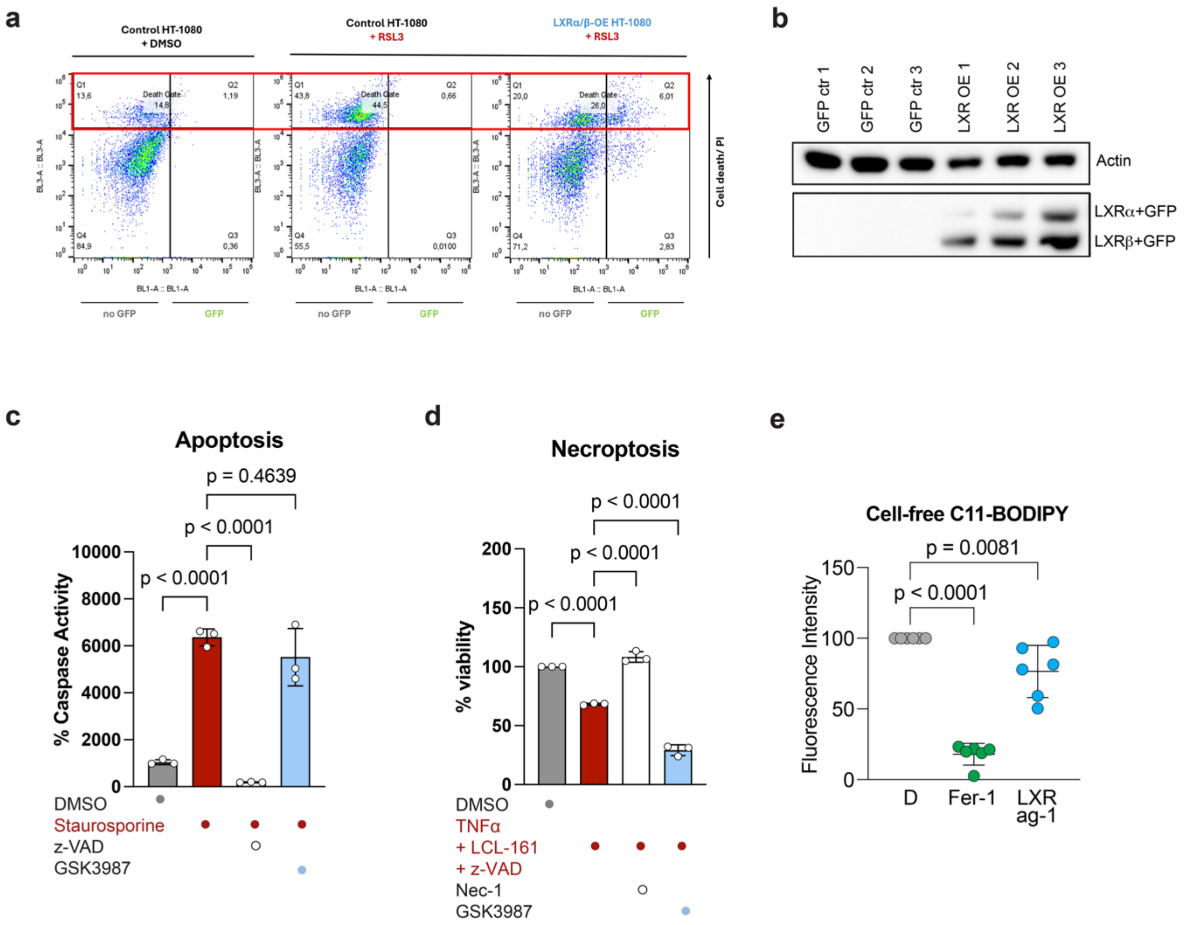


**Supplementary Figure 1: LXR overexpression and activation protect from ferroptosis**

**a)** Gating strategy for cell death analysis based on PI-positive signals following induction of ferroptosis with 200 nM RSL3. LXR-overexpressing HT-1080 cells show reduced cell death compared to control-transfected cells, as shown by the decreased proportion of PI-positive cells within the "Death Gate," including both GFP-negative and GFP-positive populations. **b)** Western blots show upregulation of LXRα and LXRβ isoforms in LXR-overexpressing HT-1080 cells compared to control-transfected cells. Protein expression of β-actin was used as a control. **c)** Treatment with 10 µM LXRag1 (GSK3987) did not reduce caspase activity in HT-1080 cells, indicating that LXR activation does not inhibit apoptosis. Staurosporine (1 µM) and z-VAD-FMK (50 µM) were used as apoptosis inducer and inhibitor, respectively; mean ± SD of *n* = 3 biological replicates; ordinary one-way ANOVA with Šidák's multiple-comparison test. **d)** Treatment with 10 µM LXRag1 (GSK3987) did not reduce the loss of viability induced by necroptosis induced with TNFα (20 ng/mL), LCL-161 (10 µM), and z-VAD-FMK (10 µM). Nec-1 was used as necroptosis inhibitor; mean ± SD of *n* = 3 biological replicates; ordinary one-way ANOVA with Šidák's multiple-comparison test. **e)** Cell-free oxidizable BODIPY-C11 assay treated with 7.5 mM free-radical-producing 2,2′-azobis(2-methyl-propanimidamide) dihydrochloride (AAPH) indicates no antioxidant capacity of LXRag1 (10 µM); mean ± SD of *n* = 3 biological replicates; ordinary one-way ANOVA with Šidák's multiple-comparison test.

#### Supplementary Figure 2

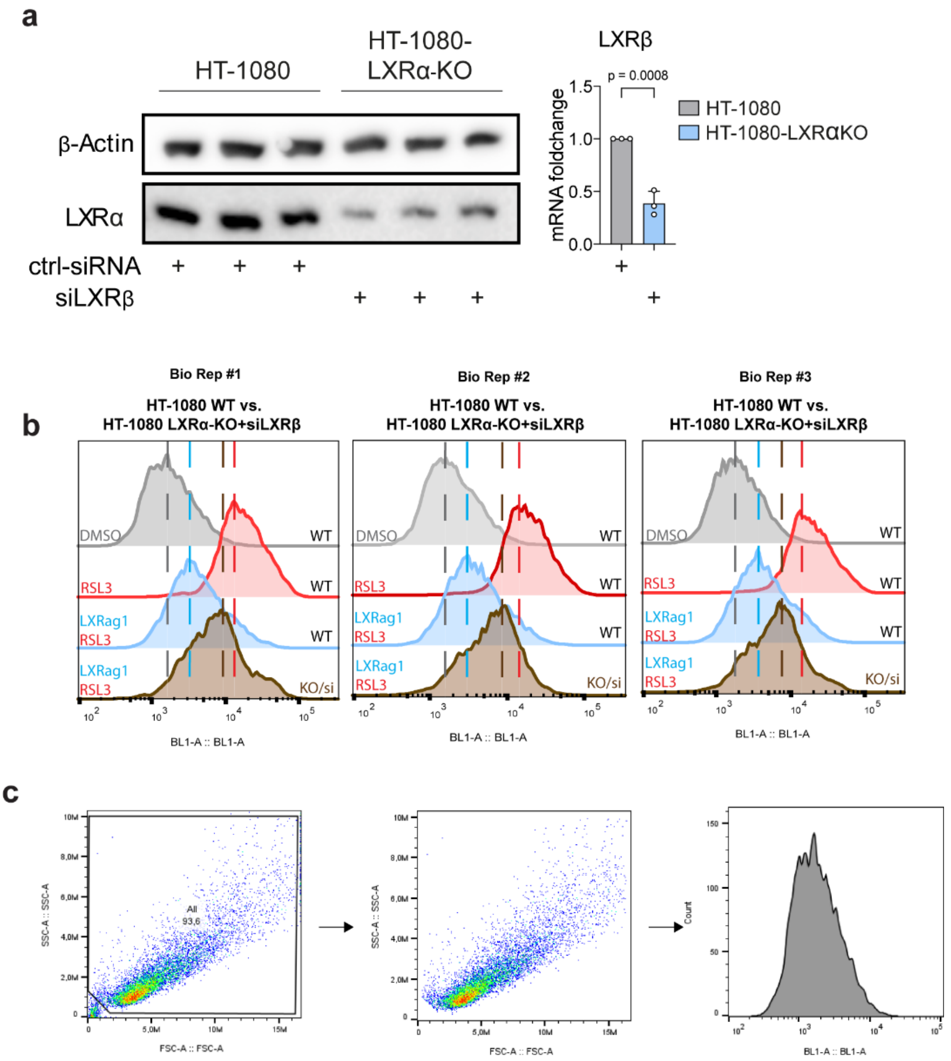


**Supplementary Figure 2: LXR knockout sensitizes towards ferroptosis**

**a)** Western blot analysis shows approximately 80% loss of LXRα protein expression in HT-1080 LXRα-KO cells; protein expression of β-actin serves as control. qRT-PCR analysis shows approximately 60% knockdown of LXRβ expression in siLXRβ-transfected cells compared to control-transfected cells; mean ± SD of *n* = 3 biological replicates; unpaired t-test. **b)** Analysis of C11-BODIPY fluorescence shift indicates that activation of LXR (10 µM LXRag1) decreases lipid peroxidation in ferroptotic (1 µM RSL3) wild-type HT-1080 cells. This reduction is attenuated in 1 µM RSL3-treated LXRα-knockout + LXRβ siRNA-transfected cells; depicted are 3 biological replicates. **c)** Gating strategy for C11-BODIPY fluorescence analysis of LXRα-knockout + LXRβ siRNA-transfected cells treated with 1 µM RSL3 and control transfected cells.

#### Supplementary Figure 3

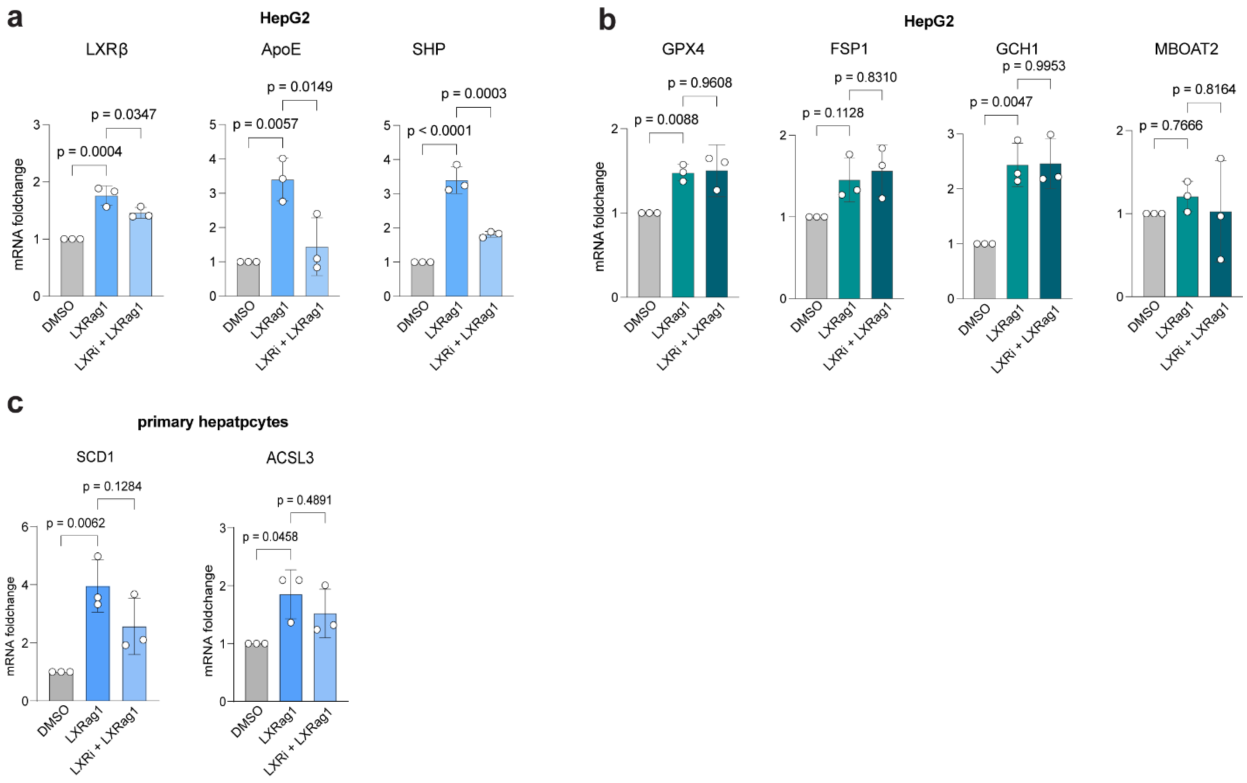


**Supplementary Figure 3: LXR promotes expression of genes involved in MUFA-PL synthesis**

**a)** qRT-PCR measurements of LXRb, ApoE and SHP show increased mRNA foldchange upon LXR activation (10 µM LXRag1) and decrease upon co-treatment with LXRi (25 µM) in HepG2; mean ± SD of *n* = 3 biological replicates; ordinary one-way ANOVA with Šidák's multiple-comparison test. **b)** qRT-PCR measurements of GPX4, FSP1 and GCH1 show slight increase in mRNA foldchange upon LXR activation (10 µM LXRag1) that does not decrease upon co-treatment with LXRi (25 µM) in HepG2. MBOAT2 shows no regulation in mRNA foldchange upon LXR activation (10 µM LXRag1) or co-treatment with LXR inhibitor (LXRi 25 µM); mean ± SD of *n* = 3 biological replicates; ordinary one-way ANOVA with Šidák's multiple-comparison test. **c)** qRT-PCR measurements of SCD1 and ACSL3 show increased mRNA foldchange upon LXR activation (10 µM LXRag1) and decrease upon co-treatment with LXRi (25 µM) in murine primary hepatocytes; mean ± SD of *n* = 3 biological replicates; ordinary one-way ANOVA with Šidák's multiple-comparison test.

#### Supplementary Figure 4

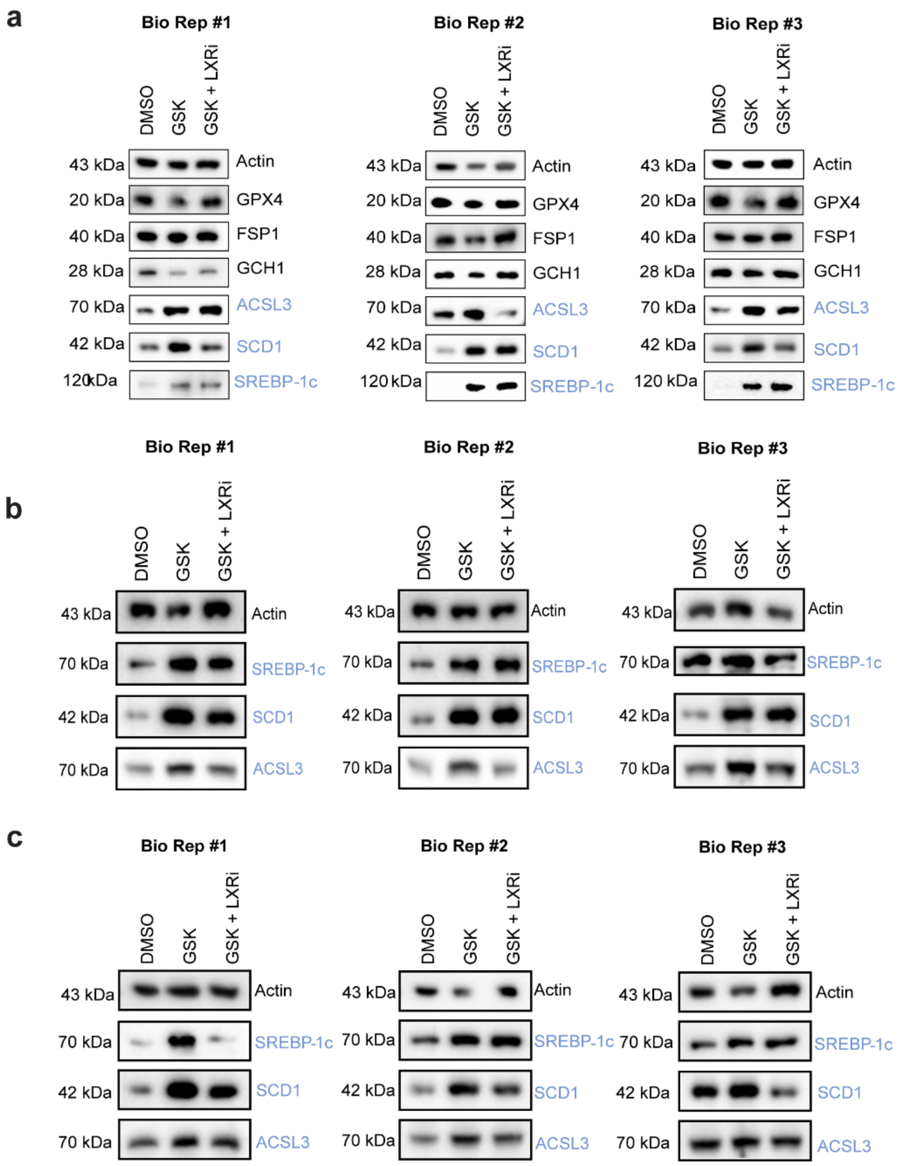


**Supplementary Figure 4: LXR promotes expression of proteins involved in MUFA-PL synthesis**
**a)** Western Blot analysis of HepG2 cells treated with LXRag1 (10 µM) with or without co-treatment with LXRi (25 µM) shows no regulation of protein levels of GPX4, FSP1 and GCH1. Protein levels of ACSL3, SREBP-1c and SCD1 show an increase upon LXR activation (LXRag1 10 µM), which is reduced upon co-treatment with LXRi (25 µM); depicted are 3 biological replicates; protein expression of β-actin serves as control. **b, c)** Western Blot analysis of HT1080 (b) and HT-29 (c) show increased protein levels of SREBP-1c, SCD1 and ACSL3 upon LXR activation (LXRag1 10 µM), which is reduced by co-treatment with LXRi (25 µM); depicted are 3 biological replicates; protein expression of β-actin serves as control.

### Supplementary Figure 5

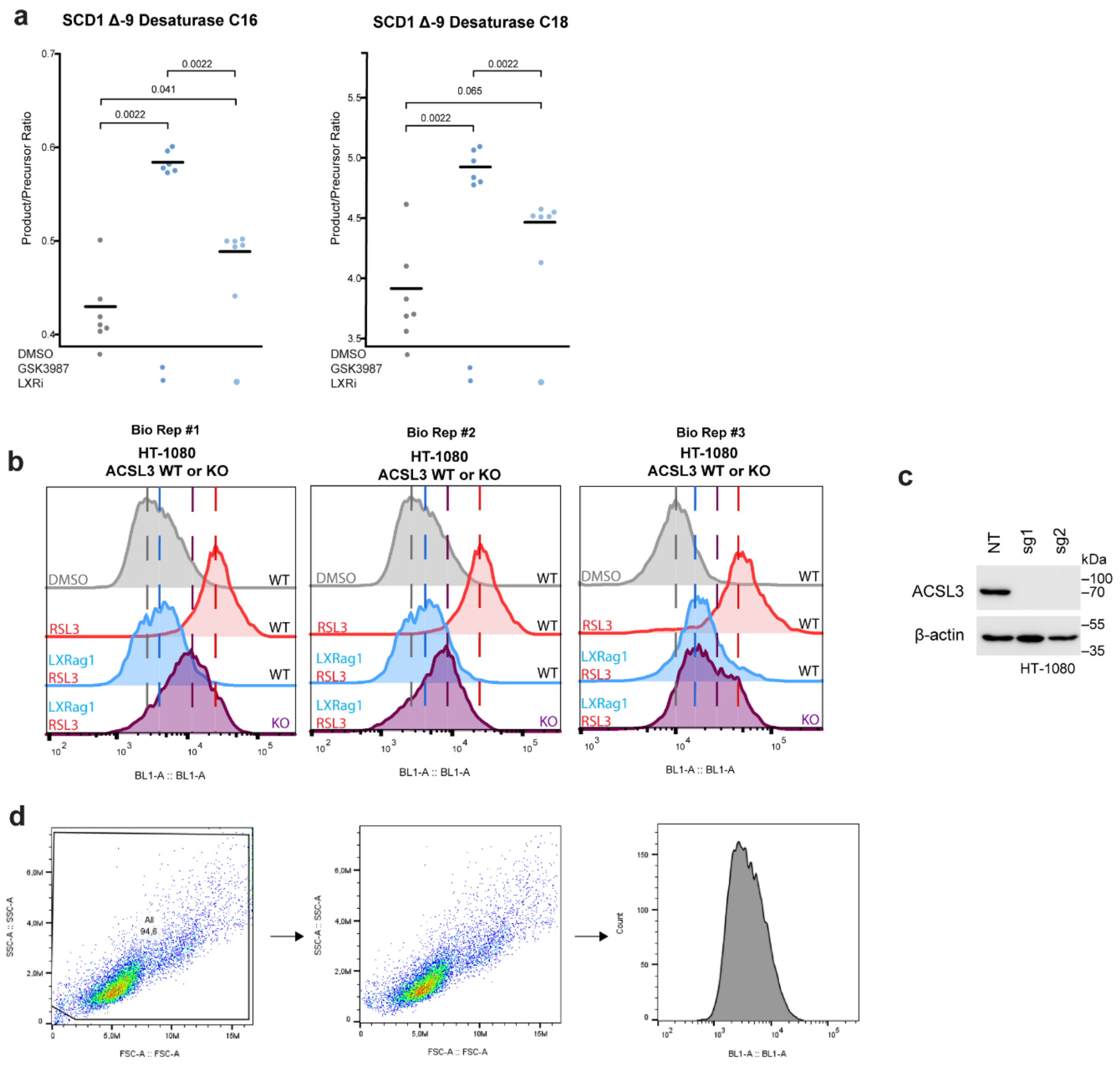


**Supplementary Figure 5: LXR protects against ferroptosis via MUFA-rich membrane remodeling**
**a)** Lipidomics-derived product-to-precursor ratios indicate increased SCD1 activity, reflected by elevated D9D(C16) and D9D(C18) ratios following LXR activation (10 μM GSK3987). Co-treatment with LXRi (25 μM) reduced this effect; mean ± SD of *n* = 6 biological replicates; unpaired Wilcoxon test. **b)** Analysis of C11-BODIPY fluorescence shift indicates that activation of LXR (10 μM LXRag1) decreases lipid peroxidation in ferroptotic (1 μM RSL3) ACSL3 wild-type HT-1080 cells. This reduction is attenuated in 1 μM RSL3-treated HT1080 ACSL3-knockout cells; depicted are 3 biological replicates; **c)** Western Blot analysis of HT-1080 ACSL3-KO shows reduced protein levels of ACSL3 compared to control cells (NT), protein expression of β-actin serves as control. **d)** Gating strategy for C11-BODIPY fluorescence analysis of HT1080 ACSL3-KO cells treated with 1 μM RSL3 and ACSL3 wild-type control cells.

#### Supplementary Figure 6

a

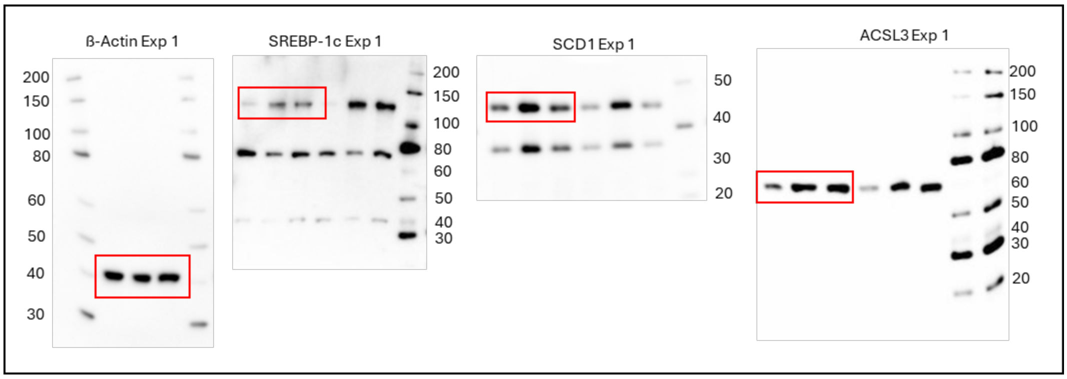


b

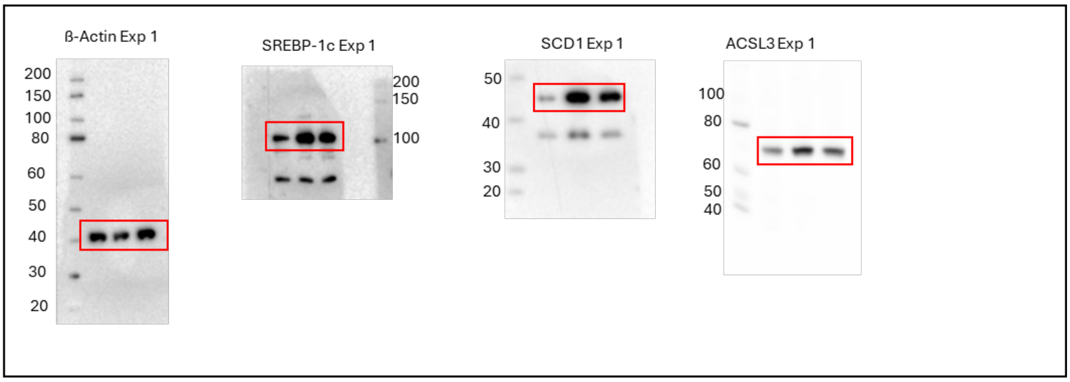


c

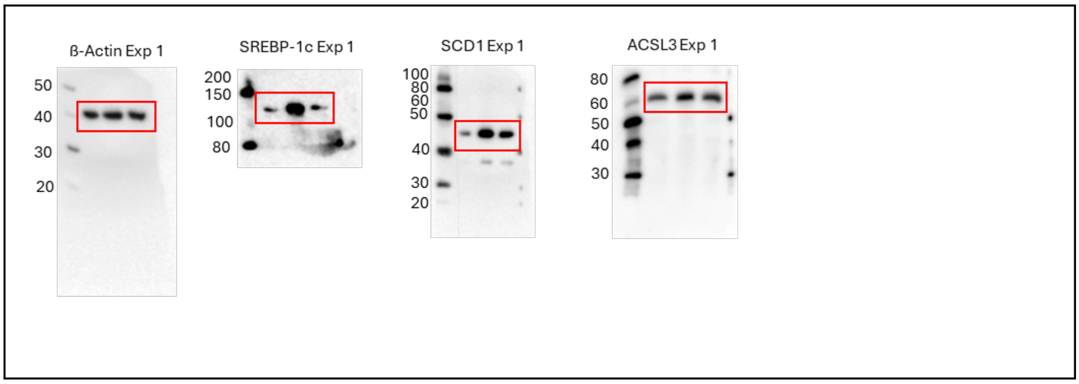


**Supplementary Figure 6: Western Blot analysis of LXR activation**
**a, b, c)** Uncropped Western Blots corresponding to Figure 4d for HepG2 (a), HT-1080 (b) and HT-29 (c) with staining against β-Actin, SREBP-1c, SCD1 and ACSL3 respectively; the WesternFroxx all-in-one Protein Ladder (neoFroxx) was used.

#### Supplementary Figure 7

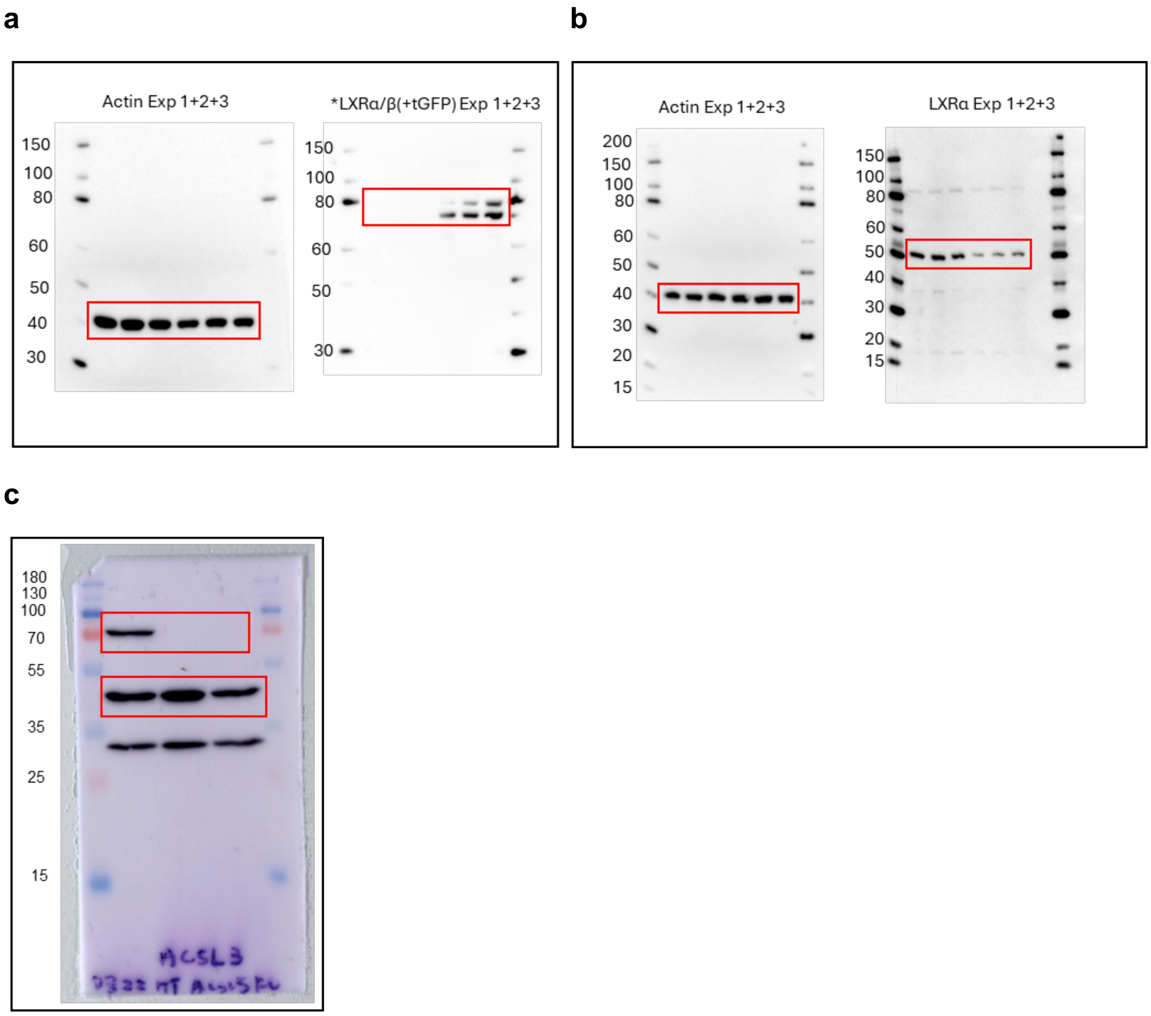


**Supplementary Figure 7: Western Blot analysis of LXR overexpression and LXRα and ACSL3 knockout**
**a)** Uncropped Western Blots corresponding to Figure 1e for HT-1080 LXRα/β-OE cells compared to control cells with staining against β-Actin and LXRα/LXRβ; the WesternFroxx all-in-one Protein Ladder (neoFroxx) was used. **b)** Uncropped Western Blots corresponding to Figure 1j for HT-1080 LXRαKO+siLXRβ cells compared to control cells with staining against β-Actin and LXRα; the WesternFroxx all-in-one Protein Ladder (neoFroxx) was used. c) Uncropped Western Blots corresponding to Suppl. Figure 2 for HT-1080 ACSL3-KO cells compared to control cells with staining against β-Actin and ACSL3, PageRuler Plus Prestained Protein Ladder was used (Thermo Fisher Scientific).

#### Supplementary Figure 8

a

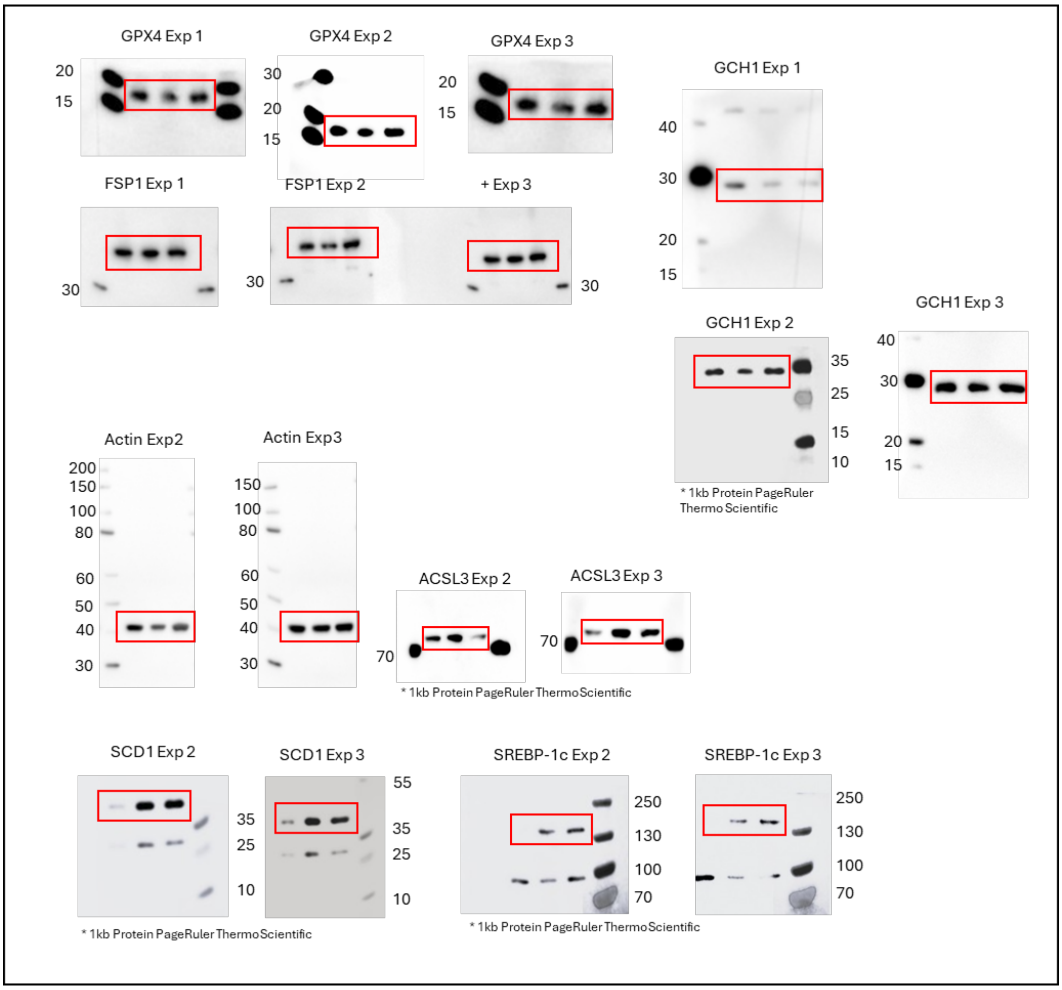


b

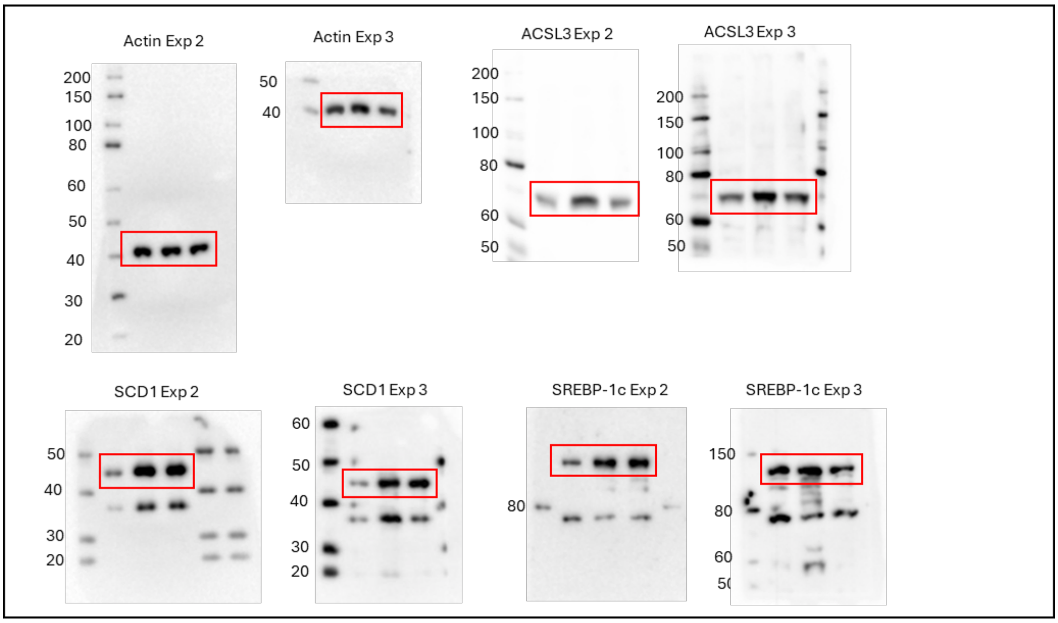

**c**

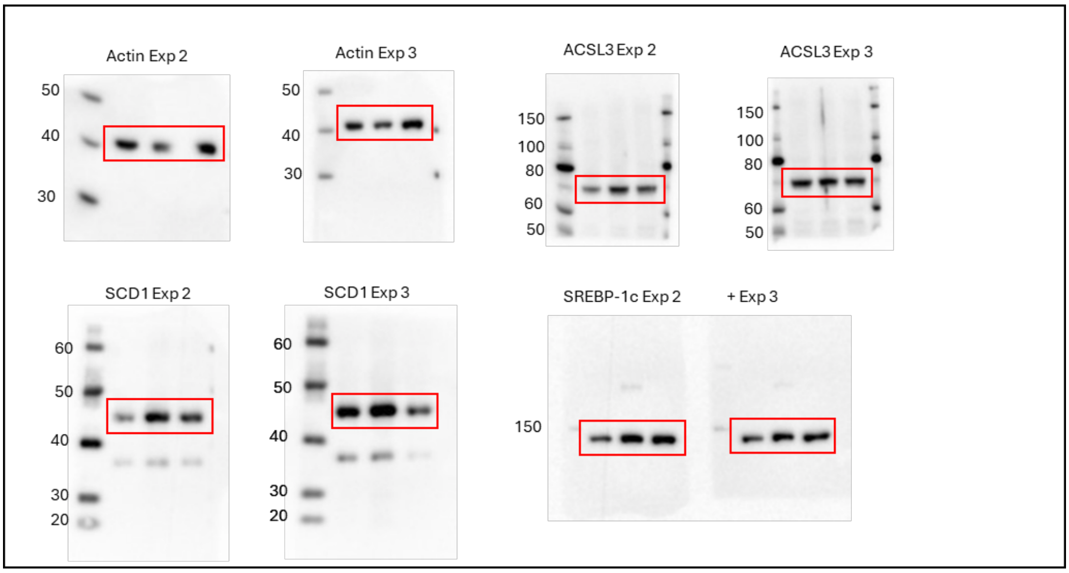


**Supplementary Figure 8: Replicates of Western Blot analysis of LXR activation**

**a, b, c)** Uncropped Western Blots corresponding to Supp. Figure 4 of HepG2 (a) with staining against , with staining against β-Actin, GPX4, GCH1, FSP1, SREBP-1c, SCD1 and ACSL3 and HT-1080 (b) and HT-29 (c) with staining against β-Actin, SREBP-1c, SCD1 and ACSL3 respectively; the WesternFroxx all-in-one Protein Ladder (neoFroxx) was used.
